# A shade-responsive microprotein in *ATHB2* regulates Arabidopsis growth and root development

**DOI:** 10.1101/2024.02.01.578400

**Authors:** Ashleigh Edwards, Maurizio Junior Chiurazzi, Anko Blaakmeer, Ylenia Vittozzi, Anne van Humbeeck, Krishna Kodappulli Das, Shaochun Zhu, Andre Mateus, Ashish Sharma, Sanne Matton, Valdeko Kruusvee, Daniel Straub, Giovanna Sessa, Monica Carabelli, Giorgio Morelli, Stephan Wenkel

## Abstract

Shade triggers widespread changes in transcript isoform production in *Arabidopsis thaliana*. Using 5’ PEAT sequencing, we identified 377 candidate microprotein-encoding transcripts arising from alternative transcription start sites, including one located within the third exon of ATHB2, a class II HD-ZIP transcription factor central to shade avoidance. The encoded microprotein, ATHB2miP, lacks the DNA-binding homeodomain but retains the leucine zipper domain, through which it forms heterodimers with full-length ATHB2, inhibits its DNA-binding activity, and disrupts its localization to nuclear photobodies. Ectopic expression of ATHB2miP phenocopies loss of ATHB2, deregulating genes involved in auxin signaling, root development, and iron homeostasis. Complementation of the *athb2* mutant with a construct carrying a silent mutation at the ATHB2miP start codon results in exaggerated hypocotyl elongation under both white light and shade, demonstrating that ATHB2miP is required to restrain excessive elongation growth. Immunoprecipitation mass spectrometry confirmed that this mutation redirects translation to alternative initiation sites, producing distinct proteoforms absent from wild-type plants. Additionally, ATHB2 suppresses iron uptake genes, and iron availability modulates shade growth responses in an ATHB2-dependent manner. Together, these findings establish ATHB2miP as a regulator of an ATHB2 feedback circuit that integrates light and nutritional signals to coordinate shoot elongation and root development.

## Introduction

Since the proposition of the “one gene - one enzyme hypothesis” by George W. Beadle and Edward L. Tatum in 1941 (Beadle & Tatum, 1941), it has become increasingly evident that the complexity of the proteome is more intricate than initially anticipated. It is firmly established that alternative splicing can give rise to multiple mRNA isoforms (Berget *et al*, 1977), and in certain instances, these isoforms can be translated into different protein variants. Alongside alternative splicing events, such as exon-skipping, which can potentially result in protein isoforms with distinct structures, the alternative use of transcription start sites can also lead to protein variants lacking specific domains that exhibit contrasting functions compared to the full-length variant (Svoboda *et al*, 1988).

Recent endeavors in uncovering the small peptidome, employing riboproteomics, a combination of Ribo-seq and proteomics, have unveiled thousands of novel protein-coding genes in humans (Mudge *et al*, 2022). Some of these short open reading frames (sORFs) originate from their corresponding loci in an overlapping manner and are frequently encoded in an alternate reading frame. Certain upstream sORFs can possess a regulatory role in controlling the larger downstream ORF at the transcriptional or translational level (Luo *et al*, 1995; Orbach *et al*, 1990). Furthermore, long non-coding RNAs (lncRNAs), which arise from pervasive transcription of the seemingly non-coding regions of the genome, represent another abundant source of small and potentially emerging protein-coding genes. It has been suggested that translated lncRNAs may even serve as a source for *de novo* gene birth (Ruiz-Orera *et al*, 2020).

Small proteins are abundant in both prokaryotes and eukaryotes, and many of them exhibit dynamic expression patterns. Notably, sORFs have been demonstrated to possess significant regulatory roles in plant development (Hanada *et al*, 2013) as well as in animals (Kondo *et al*, 2010; Zhang *et al*, 2020). In the field of human biology, certain sORFs have been identified to exhibit disease-specific expression patterns, suggesting their potential utility as diagnostic markers or future therapeutic targets (Chong *et al*, 2020).

When compared to animals, plants display a greater degree of developmental plasticity and exhibit substantial changes in gene expression in response to environmental fluctuations. Due to their sessile nature, plants constantly monitor their surroundings, particularly in relation to light, which is their main source of energy. Plants possess various photoreceptor systems, such as phytochromes and cryptochromes, which enable them to continuously assess the quality of light and determine their growth conditions in competition with other plants (Halliday *et al*, 1994; Pedmale *et al*, 2016). Phytochrome photoreceptors respond to both red and far-red light in a toggle switch manner where red light activates PHYB and converts it to the far-red light-responsive Pfr form, while far-red light inactivates PHYB and converts it to the inactive Pr form (Quail, 2002). Shade-sensitive plants like Arabidopsis promote elongation growth in response to a low red-to-far-red ratio. This growth response is mediated by the stabilization of PHYTOCHROME INTERACTING FACTOR (PIF) transcription factors (Lorrain *et al*, 2008). Under conditions of high red light (no shade), PHYB exists in the active Pfr form, leading to the degradation of PIFs. When present, PIFs directly activate genes encoding YUCCA enzymes, resulting in increased production of the plant hormone auxin in response to shade (Müller-Moulé *et al*, 2016; Won *et al*, 2011).

In addition to the pivotal role played by PIFs in regulating growth in response to shade, the class II homeodomain leucine zipper (HD-ZIPII) family of transcription factors has long been recognized as being light-regulated (Carabelli *et al*, 1993). Among the 10 genes that encode HD-ZIPII transcription factors, for *HAT1*, *HAT2*, *HAT3*, *ATHB2*, and *ATHB4*, it has been demonstrated that their expression is transcriptionally upregulated in response to shade (Carabelli *et al*, 2013; Ciarbelli *et al*, 2008). The presence of an EAR-domain in all of these HD-ZIPII genes suggests their function as transcriptional repressors. This was confirmed experimentally for ATHB2, whose EAR motif was shown to directly repress the HAT1 promoter in reporter assays (Tiwari *et al*, 2012). Initial experiments showed that ATHB2 is capable of binding to a *cis*-element within its own promoter to suppress its own expression (Ohgishi *et al*, 2001; Steindler *et al*, 1999). ATHB2 was in fact the first plant homeobox gene for which negative autoregulation was demonstrated. The same study showed that ATHB2 recognition sequences are present in the promoters of other HD-ZIPII members as well, including ATHB4, HAT1, HAT2, HAT3, HAT9, and HAT22, indicating that ATHB2 acts as a broad repressor across the subfamily (Ohgishi *et al*., 2001). In addition to their role in self-regulation, HD-ZIPIIs have been found to influence the expression of genes encoding enzymes and signaling components involved in hormone pathways such as auxin, brassinosteroids, and gibberellic acid, all of which are known to promote growth (Sorin *et al*, 2009). ATHB2 is itself an early auxin-responsive gene: transcript levels rise approximately threefold within 30 minutes of auxin treatment. Conversely, ATHB2 overexpression reduces the expression of PIN auxin efflux carriers and YUCCA biosynthetic enzymes, placing ATHB2 within a feedback circuit that modulates auxin distribution and transport (He *et al*, 2020). Furthermore, HD-ZIPIIs have been shown to play a crucial role in leaf and meristem development, as loss-of-function mutants exhibit severe defects in maintaining meristems and developing leaves (Bou-Torrent *et al*, 2012; Turchi *et al*, 2015). Interestingly, REVOLUTA (REV), a member of the class III homeodomain leucine zipper (HD-ZIPIII) family, acts upstream of HD-ZIPII factors and also participates in the regulation of shade-induced growth by transcriptionally upregulating *HAT2*, *HAT3*, *ATHB2*, and *ATHB4* (Brandt *et al*, 2012). Furthermore, it has been discovered that a dimer composed of HD-ZIPII and HD-ZIPIII can bind to a conserved *cis*-element in the microRNA genes *MIR165/166*, leading to the repression of their expression specifically in the adaxial domain of developing leaves (Merelo *et al*, 2016). Studies have also revealed that shade influences the rate of leaf proliferation, which is controlled by ATHB2 and ATHB4 (Carabelli *et al*, 2018). Beyond its roles in shade-regulated growth, ATHB2 also functions as a negative regulator of light-promoted seed germination by increasing ABA sensitivity and upregulating downstream germination inhibitors including ABI3, ABI5, and PIF1 (Tognacca *et al*, 2021). Additionally, in the hypocotyl, REV has been found to regulate xylem proliferation in a shade-dependent manner, likely providing structural support to the elongating hypocotyl while increasing water flow (Botterweg-Paredes *et al*, 2020).

HD-ZIPIII factors act as key regulators of meristem maintenance and leaf development, and therefore their regulation occurs at multiple levels, including post-transcriptional and post-translational. At the post-transcriptional level, *microRNA165/166* modulates *HD-ZIPIII* expression, limiting it to the adaxial domain of developing leaves (Mallory *et al*, 2004). At the post-translational level, HD-ZIPIII factors are subject to regulation by LITTLE ZIPPER (ZPR) microproteins (Kim *et al*, 2008; Wenkel *et al*, 2007). Microproteins are short regulatory proteins consisting of approximately 100 amino acids or less and some share structural similarities with the proteins they regulate (Bhati *et al*, 2021; Eguen *et al*, 2015). The ZPR microproteins specifically possess a leucine zipper domain similar to that found in HD-ZIPIII factors, and they are directly and positively regulated by HD-ZIPIII transcription factors (Brandt *et al*, 2013). Evolutionarily, ZPRs are related to HD-ZIPIII factors, and it has been demonstrated that ZPRs originated from a duplication of an ancient HD-ZIPIII paralog in the euphyllophyte clade, followed by evolutionary trimming through the accumulation of degenerative mutations, resulting in the preservation of only the leucine zipper domain (Floyd *et al*, 2014).

Here, we investigate how shade affects the transcription landscape in Arabidopsis. We find that in response to shading alternative transcription start sites are more often employed, potentially, increasing proteome diversity. One of the alternative transcription start sites is located in the third exon of *ATHB2* and the alternative transcript encodes a microprotein resembling the ZPRs that is devoid of the homeodomain DNA binding moiety but contains the leucine zipper part, necessary to interact with HD-ZIPII proteins. Ectopic expression of ATHB2miP, the microprotein isoform, inactivates ATHB2 by engaging it in a non-productive heterodimer and thereby affects growth of the hypocotyl and the root. By using ATHB2miP as a tool, we can effectively inactivate ATHB2 functions. CRISPR-induced mutations that remove the leucine zipper domain from the *ATHB2* gene result in defects in responding to shade and additional defects in root development. The finding that ATHB2 localizes to nuclear photobodies, mediated by the leucine zipper domain, supports a fundamental role for this domain in the formation of higher-order protein aggregates and signaling. Transgenic plants that ectopically express ATHB2miP exhibit overlapping transcriptome changes with *athb2* mutant plants. Gene ontology and cluster analysis reveal significant changes in genes encoding components of auxin synthesis and signaling, as well as genes controlling root development and nutrient uptake, specifically iron. After discovering that genes related to iron uptake were upregulated in *athb2* mutants, we measured the internal iron levels in both shoots and roots. Our findings showed that the levels were increased in both *athb2* mutants and transgenic plants that ectopically expressed ATHB2miP. Experiments on shade growth conducted on low and high iron media revealed that the availability of iron affects shoot and root growth, which is dependent on the activity of ATHB2. In summary, these findings demonstrate that ATHB2 and ATHB2miP are key mediators between shoot and root growth in response to environmental fluctuations and the nutritional status of the plant.

## Results

### The *ATHB2* gene harbors an alternative transcription start site in the third exon

To mine the transcriptome of *Arabidopsis thaliana* for novel microproteins regulating shade avoidance, we used 5’ PEAT sequencing, a method that enriches capped RNA transcripts and results in reads that correspond to transcription start sites (Ni *et al*, 2010). Plants were grown for ten days in a white light regime (WL; R:FR = 7.7) and then exposed to additional far-red light, to simulate shade (WL+FR; R:FR = 0.13; deep shade). After shade treatment for 45-and 90-minutes samples, along with an untreated control (0 minutes shade) were collected. Overall, we identified 12,398 transcription start sites (TSSs). The majority of these corresponded to putative TSSs from TAIR10, but 40% (4,982) corresponded to sites that had not previously been annotated (Fig. 1A). Most of the reads mapped to the 5’UTR of already annotated genes, but a subset was found to map to regions within gene bodies. As we were particularly interested in identifying novel transcripts that might encode microproteins (i.e. overlapping gene products), we focused on the subset of genes encoding such alternative microproteins. We therefore established a set of criteria in order to identify sites in our dataset that might correspond to an alternative microprotein mRNA transcript. Firstly, we limited our search to open reading frames (ORFs) that would encode a protein within the size range of a typical microprotein of 5-15 kDa but would differ significantly in size to the overlapping full-length gene product. Secondly, we selected for ORFs that contained a single protein-protein interaction domain allowing the alternative proteins to interfere with the full-length isoforms by forming heterodimers. Finally, we considered the biological relevance of each alternative transcript: the number of reads must exceed a certain threshold to ensure robust expression of the transcript isoform. By applying these criteria, we identified 377 potential microprotein candidates, corresponding to 3% of the total number of TSSs identified in our dataset (Fig.1A, Supplementary dataset 1).

**Figure 1.**
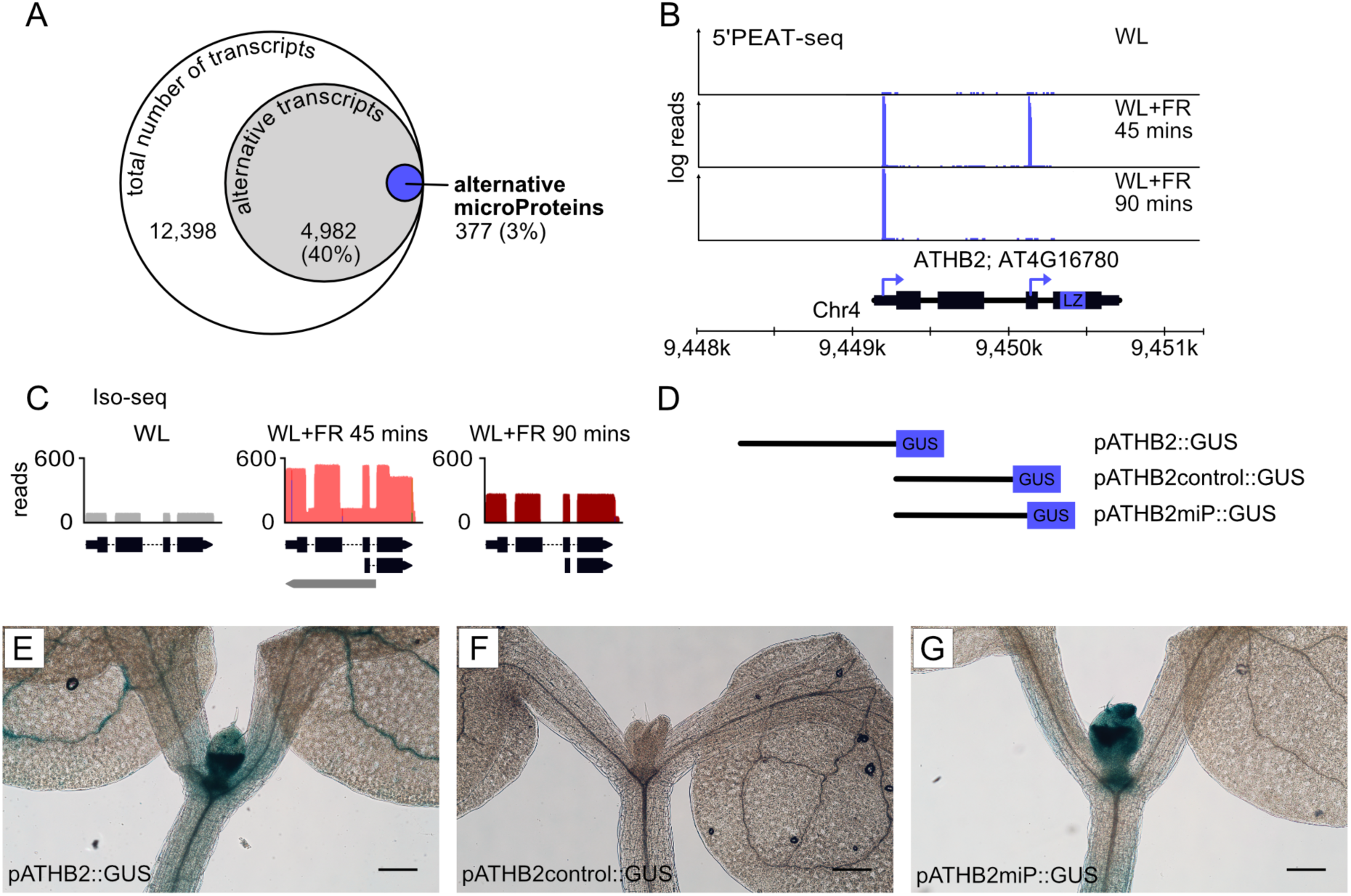
Alternative transcription from the third exon of *ATHB2* generates a shade-responsive transcript with microprotein-coding potential. **(A)** 5’PEAT-seq reads map to 12,398 unique TSSs, 43% representing novel TSSs and 1.5% being candidates for alternative microproteins (Supplementary dataset 1). **(B)** 5’PEAT-seq reads mapping to *ATHB2*. WL = White Light; FR = Far-red light. **(C)** Iso-seq reads mapping to *ATHB2*. **(D)** Design of constructs with GUS fused to either the promoter of *ATHB2* (*pATHB2::GUS*), the sequence upstream of an ATG in the second intron of *ATHB2* (*pATHB2control::GUS*; negative control), or the sequence upstream of a near-cognate start codon TTG in the third exon of *ATHB2* (*pATHB2miP::GUS*). **(E)** GUS staining in *pATHB2::GUS* accumulates in the hypocotyl, SAM and cotyledon vasculature. **(F)** GUS staining in *pATHB2control::GUS* does not produce any GUS signal. **(G)** GUS staining in *pATHB2miP::GUS* plants accumulates in the SAM and hypocotyl. Scale bars = 200 µM. All samples (E-G) had been subjected to deep shade treatment prior sampling.

One of the alternative microprotein candidates we identified overlaps with the end of *ATHB2* (Fig. 1B), a gene encoding a class II HD-ZIP transcription factor that plays a central role in the shade avoidance response (Ciarbelli *et al*., 2008; Schena *et al*, 1993). Like the full-length transcript, the alternative transcript appears to be shade-sensitive, with 5’PEAT-seq showing that expression peaks at 45 minutes. The *ATHB2* alternative transcript was also identified by 5’RACE (Fig. S1A) and by sequencing the PCR product, we found that the alternative transcript is spliced at the same site as the full-length transcript and starts around position 653 of the full-length transcript (Fig. S1B). In Arabidopsis, the average length of a 5’UTR is 155 bp (Srivastava *et al*, 2018), therefore, we searched for alternative start codons no more than 300 bp downstream of the alternative TSS. We identified two near-cognate start codons with strong Kozak sequence similarity (≥ 0.7 and < 0.8) using the online tool TISpredictor (Gleason *et al*, 2022). The two start codons are TTG and CTG, both in-frame with the full-length ATHB2 protein sequence and producing ORFs that are 126 and 102 amino acids long, respectively, and consisting of the leucine zipper domain, and, in the case of the former, parts of the homeodomain (Fig. S1B). It is unclear whether the alternative transcript we have identified is the result of an independent transcription event or due to post-transcriptional processing of the full-length *ATHB2* transcript involving downstream recapping (Ni *et al*., 2010). To confirm that ATHB2miP is the result of an independent transcription event, we sequenced full-length cDNAs using PacBio isoseq. This analysis revealed isoforms that start in the third exon of ATHB2, supporting an independent transcription event (Fig. 1C). Furthermore, we also observed antisense transcripts covering the introns. These changes occurred for both isoseq and the 5’PEAT-seq only at the 45-minute timepoint, indicating major chromatin reorganization in response to short-term shading. At 90 minutes, the situation stabilized, and ATHB2 was expressed at higher levels (Fig. 1C). To determine whether there are sequences in the gene body of *ATHB2* that can promote transcription and translation, we fused a β-glucuronidase *uidA* (GUS) reporter gene to different sequences from the *ATHB2* gene body (Fig. 1D). First, we fused GUS to the sequence 1 kb upstream of the *ATHB2* full-length start codon ATG (*pATHB2::GUS*). As a control, we fused GUS to the sequence immediately upstream of an ATG located in the middle of intron two (*pATHB2control::GUS*). Finally, we fused GUS to the sequence immediately upstream of the first alternative, in-frame codon that we had identified, TTG (*pATHB2miP::GUS*). Whilst *pATHB2control::GUS* did not display any GUS staining upon treatment with the substrate 5-bromo-4-chloro-3-indolyl ß-d-glucuronide (X-Gluc; Fig. 1F), both *pATHB2::GUS* and *pATHB2miP::GUS* showed staining in the shoot apical meristem (SAM) and the hypocotyl, though to a lesser extent in the latter in *pATHB2miP::GUS* (Fig. 1E, G). This suggests that there are sequences within the second intron and third exon, that are excluded from *pATHB2control::GUS*, that can promote both transcription and translation. This might also lend support to our hypothesis that the alternative transcript arises through an independent transcription event to the full-length *ATHB2* transcript.

### ATHB2miP heterodimerizes with ATHB2 to inhibit its DNA-binding activity

We hypothesized that the ATHB2miP might inhibit the full-length ATHB2 in a manner similar to that of the LITTLE ZIPPERs (Kim *et al*., 2008; Wenkel *et al*., 2007). To test this hypothesis, we posed the following questions. Firstly, does the microprotein localize to the same cellular compartment as the full-length protein? It was previously reported that ATHB4, another member of the HD ZIPII family, contains a nuclear localization signal (NLS) in its homeodomain that is highly conserved in ATHB2 (Gallemi *et al*, 2017). However, our own analysis using LOCALIZER (Sperschneider *et al*, 2017), which has been trained specifically on plant proteins, suggests that there is an NLS in the LZ domain that is conserved in other HD-ZIPIIs, including ATHB4. Interestingly, LOCALISER also predicts a chloroplast transit peptide (cTP) at residues 1-46 of ATHB2 with a probability of 0.8. The cTP does not appear to be conserved, suggesting that, if real, it is specific to ATHB2. To determine the subcellular localization of ATHB2 and ATHB2miP, we transiently expressed eGFP fusions of the proteins in *Nicotiana benthamiana* leaves (Fig. 2A; eGFP-ATHB2, eGFP-ATHB2miP). eGFP-ATHB2miP encompasses the entirety of exon 3 and 4 of ATHB2. To further test whether the homeodomain of ATHB2 could target the nucleus, as shown previously for ATHB4 (Gallemi *et al*., 2017), we also fused a variant of ATHB2 lacking the leucine zipper to eGFP (eGFP-ATHB2HD). We observed that all variants of ATHB2 localized exclusively to the nucleus, unlike the negative control (eGFP) which was dispersed along the cell membrane and in the nucleus (Fig. 2A). We also observed that the distribution of eGFP-ATHB2 was not uniform in the nucleus, but instead localized to distinct nuclear speckles or, in some case, to the nucleolus specifically. Nuclear speckles were mostly absent in leaves expressing eGFP-ATHB2miP, but entirely so in leaves expressing eGFP-ATHB2HD, suggesting that interactions mediated by the leucine zipper domain and in part by the N-terminal domain are required for the speckles to form. We next asked whether we could find evidence that ATHB2 and ATHB2miP can physically interact with each other. It has been demonstrated that ATHB2 homodimerizes in order to bind to its target DNA sequence and that this interaction is directed by the leucine zipper domain (Sessa *et al*, 1993). To test whether ATHB2 and ATHB2miP can interact, we performed a yeast-2-hybrid (Y2H) study. As in the localization assay, ATHB2 or ATHB2miP were fused to either the activation or the DNA binding domain of GAL4. Constructs were transformed into yeast carrying the *HIS3* gene fused to the target DNA sequence of *GAL4*. Growth of transformed cells on media lacking histidine revealed that ATHB2 and ATHB2miP can interact and form heterodimeric complexes as well as homodimers (Fig. 2B). To further confirm this interaction *in planta* we carried out a FRET-FLIM assay. Here again, ATHB2 and ATHB2miP were fused to either eGFP or mCherry and then transiently co-expressed in tobacco leaves. Using a confocal microscope, fluorescence lifetime of eGFP-ATHB2 was measured. We observed that in cells that co-expressed fluorescently tagged versions of ATHB2 or ATHB2miP there was a significant decrease in fluorescence lifetime of eGFP-ATHB2 when compared to cells that expressed eGFP-ATHB2 alone or with a control (mCherry; Fig. 2C,D), supporting that ATHB2 and ATHB2miP can physically interact.

**Figure 2.**
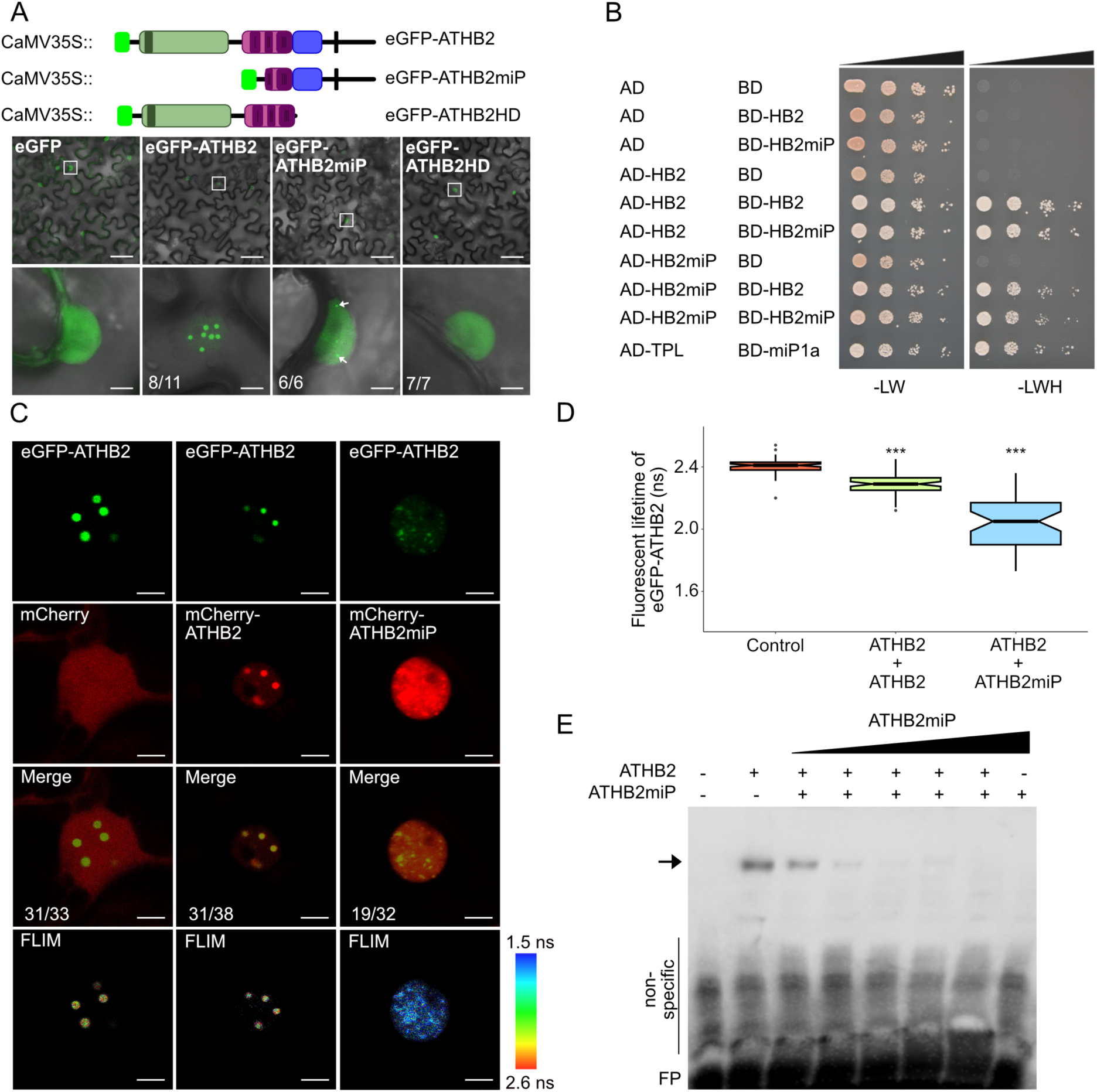
ATHB2 and ATHB2miP localise to the nucleus and interact in vitro and *in planta*. **(A)** Graphic representation of eGFP fused to ATHB2 variants: ATHB2, ATHB2miP, and ATHB2HD; panels show subcellular localisation of ATHB2 variants when fused to eGFP in tobacco leaves, with nuclear localisation shown in square and enlarged in lower panel. Arrows point to nuclear speckles that form when eGFP-ATHB2miP is expressed. Scale bars: upper panel = 50 µM; lower panel = 5 µM. Numbers indicate the proportion of nuclei displaying the depicted localisation pattern out of all nuclei examined for each construct. **(B)** Yeast-2-hybrid with growth on histidine drop-out media when ATHB2 (HB2) or ATHB2miP (HB2miP) are fused to the activation domain and binding domain of GAL4. **(C)** Co-expression of eGFP-ATHB2 with mCherry-ATHB2 or mCherry-ATHB2miP in tobacco leaves shows co-localization in the nucleus. Numbers indicate the proportion of nuclei showing co-localisation of eGFP and mCherry signals out of all nuclei examined. (**D**) FRET-FLIM assay; significant reduction in fluorescent lifetime of eGFP-ATHB2 were detected in cells co-expressing either mCherry-ATHB2 or mCherry-ATHB2miP. Scale bar = 5 µM. Significance levels: * = p<0.05, ** = p<0.01, *** = p<0.001, ns = not significant. (**E**) Electromobility shift assay demonstrating the interaction of ATHB2 with the HD-ZIPII DNA-binding site (arrow). Adding increasing amounts of ATHB2miP while keeping the levels of ATHB2 constant causes a reduction in the ability of ATHB2 to bind DNA.

Given that ATHB2 produces a full-length transcription factor and a shorter ATHB2miP isoform lacking the DNA-binding domain, we tested whether ATHB2miP interferes with ATHB2 DNA-binding activity. Recombinant ATHB2 and ATHB2miP were purified from Escherichia coli and analysed by electrophoretic mobility shift assays. ATHB2 formed a distinct complex with a labelled ATHB2 binding site, whereas addition of increasing amounts of ATHB2miP progressively reduced complex formation. These results demonstrate that ATHB2miP inhibits ATHB2 DNA binding by sequestering it into a non-DNA-binding complex (Fig. 2E).

### ATHB2miP perturbs the ability of ATHB2 to regulate transcription

To assess the functional consequences of ATHB2 inhibition by ATHB2miP, we analysed the transcriptional changes resulting from ectopic expression of ATHB2miP. For this, we grew transgenic plants overexpressing ATHB2miP, an *athb2* mutant, and wild-type plants in white light conditions. Prior to RNA extraction, plants were exposed to 0, 45 and 90 minutes of additional far-red light. RNA-seq was then used to compare the transcriptomes of respective plants under white light and shade conditions. We hypothesized that overexpressing ATBH2miP would result in the inactivation of ATHB2, similar to the inactivation of HD-ZIPIII transcription factors by ZPR microproteins (Kim *et al*., 2008; Wenkel *et al*., 2007). Differential gene expression analysis, adjusting for genotype, environment and genotype by environment interaction, identified approximately 5,900 differentially expressed genes (DEGs) with FDR ≤0.05 and significant up/down-regulation (logFC>1; logFC<-1). As it is known that HD-ZIPIIs establish an autoregulatory feedback loop by binding to their own promoter and repressing their own expression (Ciarbelli *et al*., 2008), we examined the *ATHB2* gene in all three genotypes. No reads mapping to the *ATHB2* locus were detected in the *t-athb2* mutant (Figure S2), indicating a true null allele. The transgenic plants overexpressing ATHB2miP showed an increase in the number of reads covering exons 1 and 2 compared to wild type plants, suggesting an increase in the endogenous expression of *ATHB2* (Figure S2). These findings indicate that ectopic expression of ATHB2miP sequesters the endogenous ATHB2 protein and inactivates the negative auto-repressor loop.

In total, we identified 95 DEGs whose expression depended on genotype, environment, and genotype x environment (Fig. 3A). The GO analysis of these 95 genes revealed an overrepresentation of genes involved in the shade avoidance response, growth, and hormone/auxin signaling (Fig. S3A). These findings demonstrate that both *t-athb2* mutant plants and the transgenic plants overexpressing ATBH2miP can detect shade and respond to it at the molecular level. In order to compare the genotypes and environmental conditions, we carried out an unbiased gene cluster analysis using mFuzz (Kumar & M, 2007). Our analysis revealed 17 individual clusters with distinct expression patterns (Fig. S3B). Cluster 1 (Fig. 3B) contains 200 genes that are upregulated only in *35S::miP*, but not in either of the other two genotypes. We found an overrepresentation of genes involved in root development in this cluster. Cluster 2 contains 382 genes with lower expression in both 35S::miP and *t-athb2*. This cluster is also enriched in genes encoding hormonal regulators and regulators of root development. Finally, cluster 14 contains 152 genes that are up-regulated in both *35S::miP* and *t-athb2* plants (Fig. 3B). Genes in this cluster have known roles in the shade avoidance response. We also found an overrepresentation of genes involved in iron uptake and signalling, which are strongly upregulated in *35S::miP* and *t-athb2* plants. The latter finding suggests that ATHB2 may play a role in mediating between nutritional status and growth responses.

**Figure 3.**
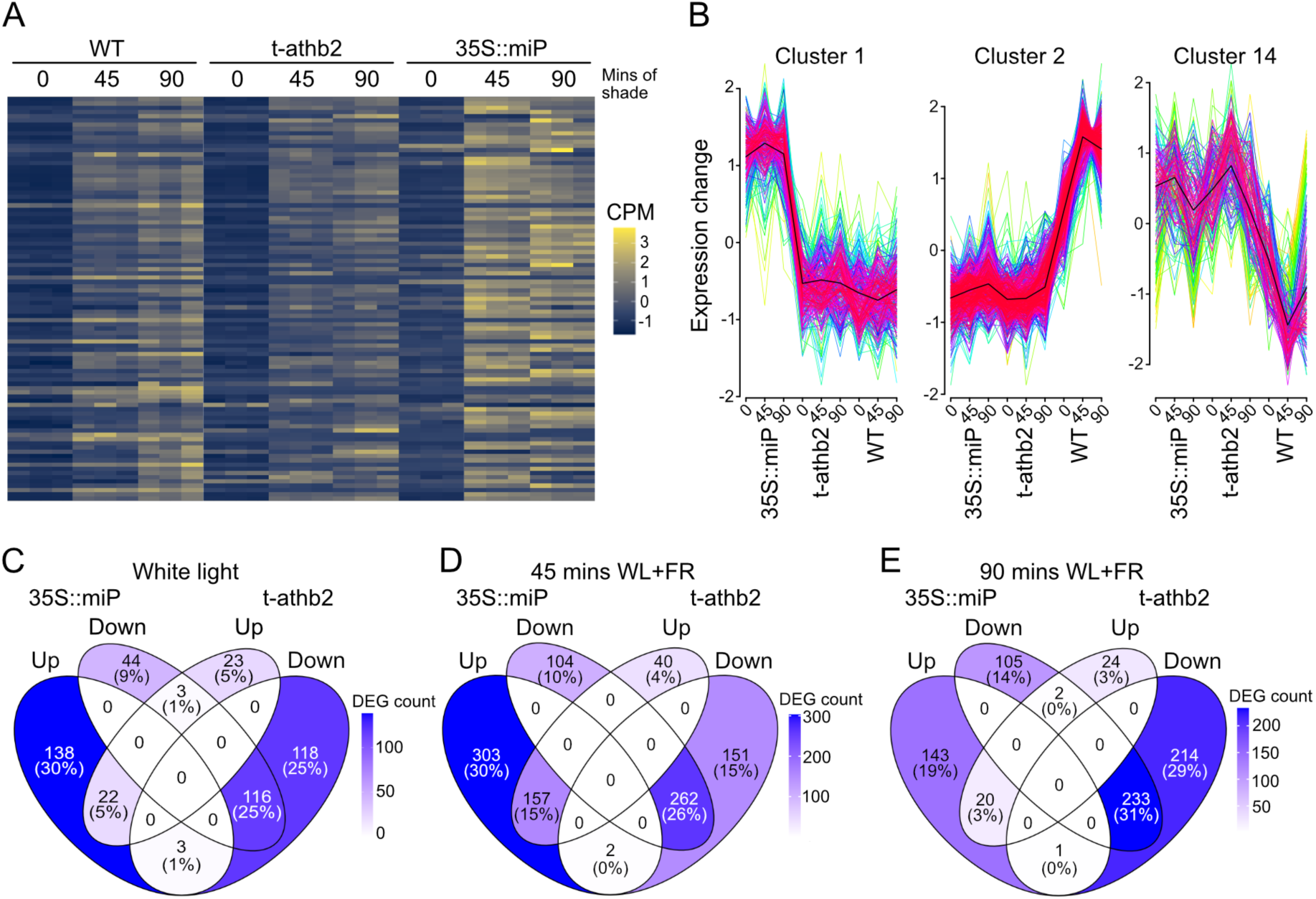
Gene expression profiling of the *t-athb2* and *35S::miP lines*. **(A)** Heatmap showing expression changes in 95 genes whose expression depends on both the genotype and the shade treatment. Lowly-expressed genes are in blue, highly-expressed genes are in yellow. **(B)** 3 of the 17 clusters generated with MFuzz. Expression changes of the genes included in each cluster are on the y-axis. The 3 genotypes and shade conditions are on the x-axis. **(C, D, E)** Venn diagrams showing the number of genes that are upregulated and downregulated in each of the three genotypes included in the RNA-Seq and the overlap between them in WL, and 45 and 90 mins of WL+FR.

Having established that ectopic expression of ATHB2miP perturbs ATHB2 function and alleviates the negative auto-repressive feedback loop we assessed DEGs in the three different environmental conditions (0-, 45-, and 90-minutes additional FR-light). Focusing on the genes upregulated in both *t-athb2* and *35S::miP* plants, we found 22 genes in white light (Fig. 3C), 157 genes in 45-minute shade (Fig. 3D), and 20 genes in 90-minute shade (Fig. 3E). As previously observed, we identified upregulated genes related to auxin synthesis and signaling, as well as genes involved in root development. Genes related to auxin-biology and hypocotyl elongation (Table 1) include *YUCCA3* and *YUCC9*, both of which have known functions in auxin biosynthesis in response to shade. (Goyal *et al*, 2016). Genes associated with root development (Table 2) include several cell wall modifying enzymes, most notably expansins, as well as transcription factors. Taken together, these findings support the function of ATHB2 as a suppressor of auxin signaling and root development. Analysis of genes down-regulated in both *t-athb2* and *35S::miP* plants showed that genes encoding defence regulators were enriched in all conditions (0, 45 and 90 min additional FR light). In addition, we found that genes involved in the biosynthesis of tryptophan were down-regulated in the shaded environments. Since auxin is mainly synthesised from tryptophan (Tao *et al*, 2008; Won *et al*., 2011), the latter finding may explain why *athb2* mutant plants have defects in shade-induced growth promotion (Ciarbelli *et al*., 2008; Schena *et al*., 1993).

**Table 1.**
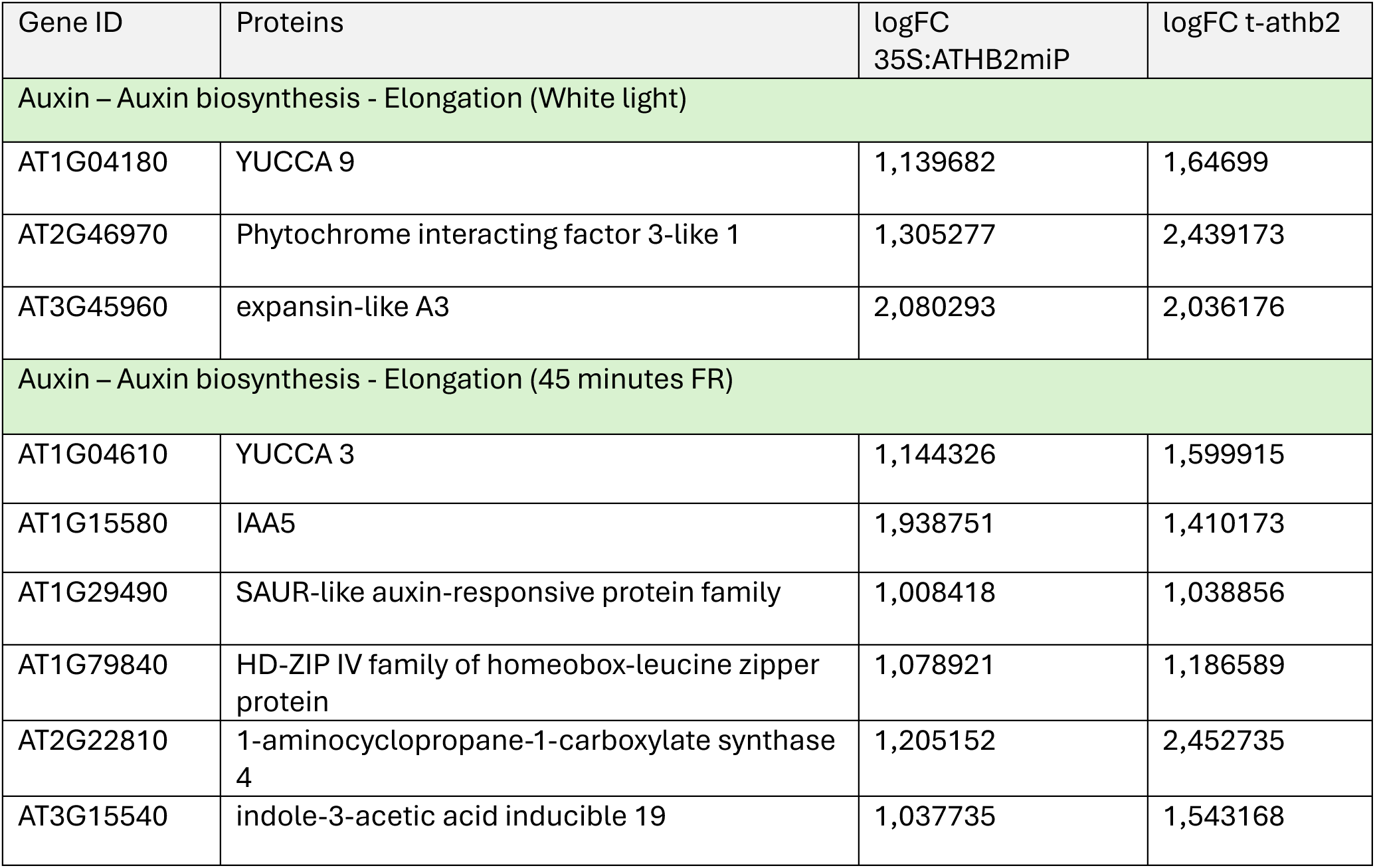
DEGs related to auxin-biology and hypocotyl elongation. Column 1: gene identifier; column 2: gene annotation; column 3: log2 fold change in 35S::ATHB2miP relative to wild type; column 4: log2 fold change in t-athb2 relative to wild type.

**Table 2.**
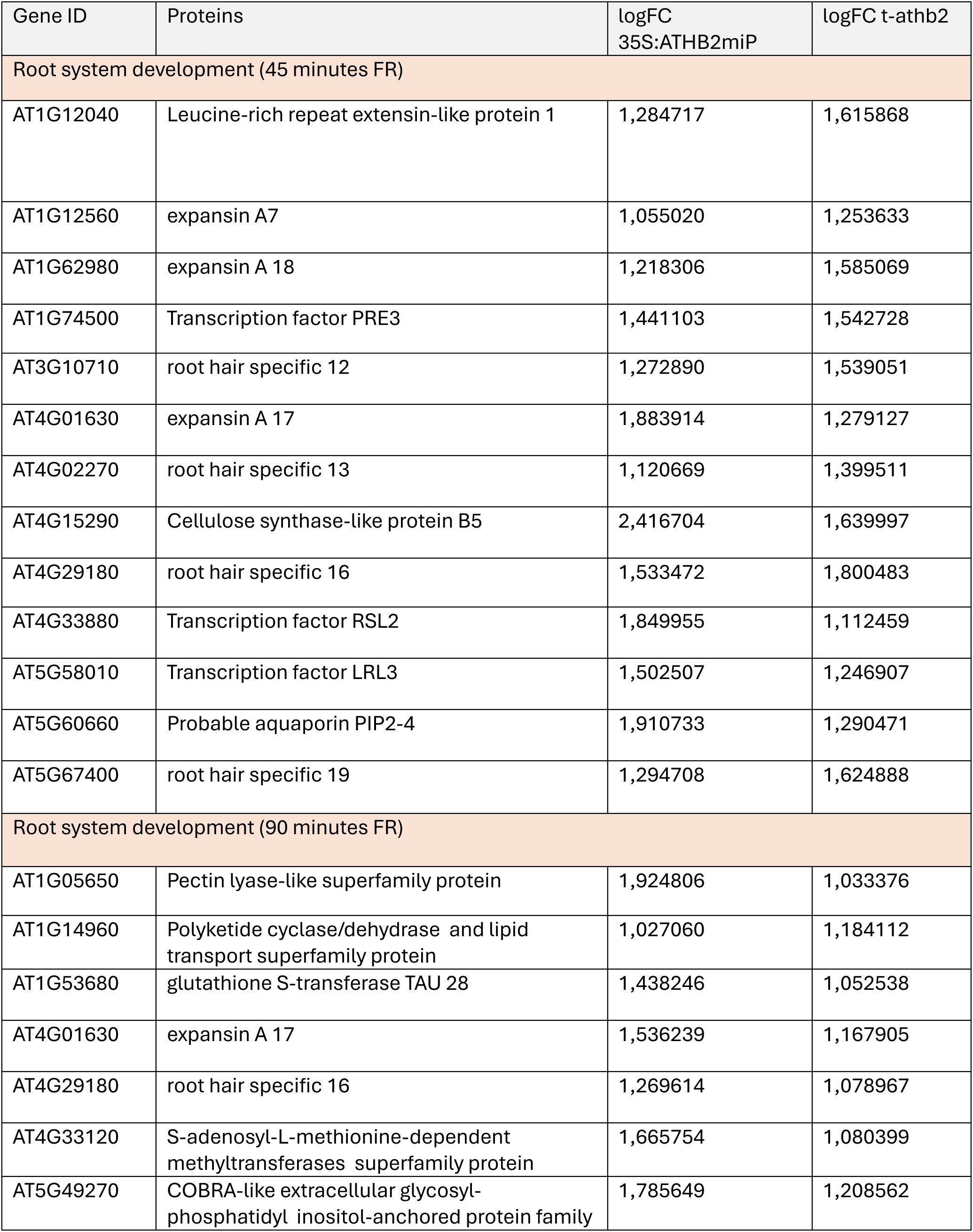
DEGs related to root development. Column 1: gene identifier; column 2: gene annotation; column 3: log2 fold change in 35S::ATHB2miP relative to wild type; column 4: log2 fold change in t-athb2 relative to wild type.

### ATHB2 controls elongation growth and root development in response to shade

In shade-avoidant plants, such as *A. thaliana*, shade caused by neighboring plants elicits a set of growth responses including elongation of stem-like structures, hyponasty and leaf extension, and acceleration of flowering (Martinez-Garcia & Rodriguez-Concepcion, 2023). It has been demonstrated that ATHB2 plays a central role in the shade avoidance response, with transgenic Arabidopsis plants where *ATHB2* is affected displaying varying degrees of shade response defects, the most obvious being in the elongation of the hypocotyl (Steindler *et al*., 1999). Early studies found that when knocked down or overexpressed, *ATHB2* was associated with subsequent shade responses where there was less or more hypocotyl elongation compared to the WT, respectively. For this reason, and because we show that the two proteins interact, we expected that transgenic plants where ATHB2miP (*35S::miP*) is overexpressed would show a phenotype indicative of lost ATHB2 function, i.e. shade insensitivity. To better understand the role of the region corresponding to ATHB2miP, we also created mutant plants by CRISPR-Cas9 gene editing to compare with the overexpression and KO line already described (Fig. 4A). The first, *athb2cr1*, presents a loss-of-function allele of *ATHB2* caused by a 25-bp deletion in the third exon, causing a frameshift and nonsense mutation, and resulting in a protein product lacking the leucine zipper and part of helix III of the homeodomain. Secondly, *athb2cr2,* is a complete loss-of-function of *ATHB2*, caused by a frameshift mutation in the beginning of the first exon, resulting in a truncated protein that shares only its first twelve amino acids with ATHB2. Based on RT-qPCR results where we analyzed expression of *ATHB2* after 45 minutes in shade, we see that transcription is not affected in any of the CRISPR lines we have generated, unlike in *t-athb2* where *ATHB2* is clearly downregulated (Fig. 4B), as also shown in the RNA-seq analysis (Fig. S2). To measure responsiveness to shade, we grew these mutant and transgenic plants alongside the Col-0 wild-type (WT) under three different shade regimes (Fig. 4C). We grew plants in deep shade (13 µmol m^-2^ s^-1^ PAR in WL and WL+FR; R:FR = 0.13 in WL+FR) with reduced PAR and a very low R/FR-ratio. In addition, we used canopy shade (45 µmol m^-2^ s^-1^ PAR in WL and WL+FR; R:FR = 0.15 in WL+FR) and proximity shade (45 µmol m^-2^ s^-1^ PAR in WL and WL+FR; R:FR = 0.06 in WL+FR) with higher PAR but different F/FR-ratios. In deep shade, we observed that *athb2* mutants were hypersensitive, with slightly elongated hypocotyls in white light conditions and, as in the case of *35S::miP*, with very long hypocotyls in shade and significantly longer hypocotyls in white light conditions. Interestingly, the CRISPR mutants *athb2cr2* and *athb2cr1* exhibited responses similar to those of wild-type plants. It is worth noting that ATHB2 is still expressed in these mutants (Fig. 4B), which suggests that ATHBmiP and other transcript or protein isoforms may still be produced. Under canopy shade, the *athb2* mutants exhibited a growth behaviour that was less sensitive to shade, resulting in reduced hypocotyl elongation in shaded conditions. However, under proximity shade, there were no significant differences in hypocotyl elongation between the wild type and *athb2* mutant plants (Fig. 4C). The results indicate that ATHB2 function depends on the shade regime. In deep shade, ATHB2 acts as a growth repressor, whereas in canopy shade it promotes elongation growth. Under proximity shade conditions, no significant differences in hypocotyl elongation were detected between wild-type and *athb2* mutant plants.

**Figure 4.**
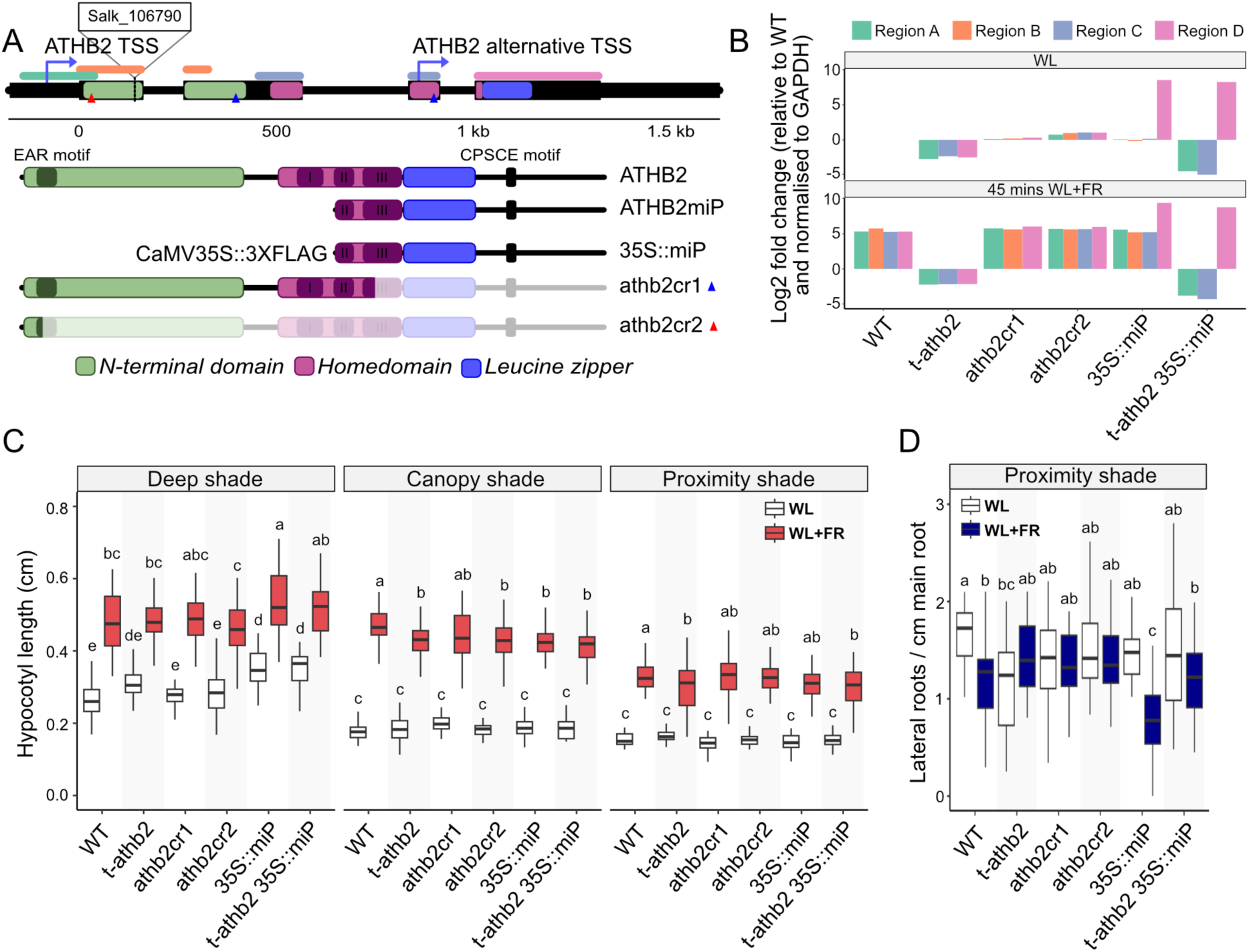
ATHB2 and ATHB2mip mediate hypocotyl elongation and root growth. **(A)** Graphic representation of the gene structure of *ATHB2*. Location of T-DNA insertion (SALK_106790) is shown and below that the predicted protein structure of WT, transgenic and mutant isoforms of *ATHB2*. Blue and red triangles indicate Cas9 targeted sites. **(B)** RT-qPCR of *ATHB2* in mutant and transgenic plants with and without WL+FR treatment. Coloured regions correspond to those in part A. **(C)** Plots of hypocotyl length in plants after 5 days growth in three different shade regimes: deep shade, canopy shade, and proximity shade. **(D)** Plot of the number of lateral roots in plants after 11 days of growth in WL or WL+FR (proximity shade). Letters in all signify results of pairwise comparisons by two-way ANOVA.

Class II HD-ZIPs have been suggested to have some redundant function in the shade avoidance response, so it is possible that ATHB2miP also interacts with these other proteins in the absence of full-length *ATHB2*. To determine whether the effect of *35S::miP* depends on the presence of the endogenous copy of *ATHB2*, we expressed *35S::miP* in the *t-athb2* knock-out. In deep shade, we observed that both *35S::miP* and *35S::miP t-athb2* plants developed longer hypocotyls under white light, indicating that ATHB2miP may have inhibited other transcription factors. Additionally, *35S::miP* plants had significantly longer hypocotyls than *35S::miP t-athb2* plants in deep shade, suggesting that the absence of full-length ATHB2 may contribute to this difference. The shade avoidance response is also associated with changes in root architecture. For example, one study found that lateral root density was decreased in plants that were exposed to longer periods of shade (van Gelderen *et al*, 2018). Mutations affecting ATHB2 have been reported to affect root development, disrupting both root length and lateral root formation (Schena *et al*., 1993), and we also identified a number of genes related to root morphogenesis in our RNA-seq analysis. Therefore, we questioned whether ATHB2miP could also have a role in root development.

Following the shade regimes described earlier, plants were grown in either WL+FR or WL for a period of 11 days and primary root length was measured. Our results indicate that there were no significant differences in all conditions tested, except for the *t-athb2* mutant plants which had slightly shorter roots in white light and even shorter roots in the proximity shade regime (Suppl. Fig. S4). We also assessed the ability of plants to form lateral roots under light and shade conditions. While no difference in the number of lateral roots respective to genotypes and environmenal conditions was observed in deep shade and canopy shade, we detected significant differences in proximity shade conditions (Fig. 4D). In white light, lateral root numbers were comparable across all genotypes. The *athb2* single mutant showed a slight reduction relative to wild type that did not reach statistical significance. Under shade conditions, *t-athb2*, a*thb2cr2*, and *athb2cr1* mutants did not differ significantly from wild type. In contrast, *35S::miP* transgenic plants showed significantly fewer lateral roots than wild type in shade. Notably, this suppression was absent in *t-athb2 35S::miP* plants, which showed lateral root numbers comparable to wild type, indicating that the effect of ATHB2miP on lateral root emergence requires functional ATHB2. Taken together, ATHB2 controlled hypocotyl elongation in a shade regime-dependent manner, acting as a growth repressor in deep shade and a growth promoter in canopy shade, a function partially masked in CRISPR alleles that may still produce ATHB2miP. The epistatic relationship between ATHB2miP and ATHB2 in lateral root regulation suggests that ATHB2miP does not suppress lateral root emergence independently, but rather acts through ATHB2, possibly by modulating its transcriptional activity in a complex whose composition or target specificity differs from that of ATHB2 homodimers.

### Loss of ATHB2miP results in a constitutive shade response under both white light and shade conditions

To genetically disentangle the functions of full-length ATHB2 and the ATHB2miP isoform, we analysed mutants lacking the entire ATHB2 protein alongside mutants in which the ATHB2miP coding region was disrupted. Because the ATHB2miP sequence is fully contained within the ATHB2 coding region, transcript-specific knockdown approaches such as RNAi cannot selectively eliminate the microprotein without also affecting the full-length proteoform. We therefore complemented the *t-athb2* mutant with a genomic *ATHB2* transgene (*ATHB2 WT*) comprising 2 kb of upstream promoter sequence including the *5’UTR*, the complete genomic locus with all exons and introns, and a C-terminal triple FLAG tag followed by 300 bp of downstream sequence encompassing the full *3’UTR*. To specifically abolish translation of ATHB2miP without altering the full-length ATHB2 protein sequence, we generated a second complementation construct (*ATHB2 alt*) in which the alternative near-cognate start codon TTG was mutated to CTC. This substitution is predicted to prevent translation initiation of ATHB2miP while preserving the encoded amino acid sequence, as both codons encode leucine. Independent homozygous transgenic lines were established in the *t-athb2* background (Fig. 5A).

**Figure 5.**
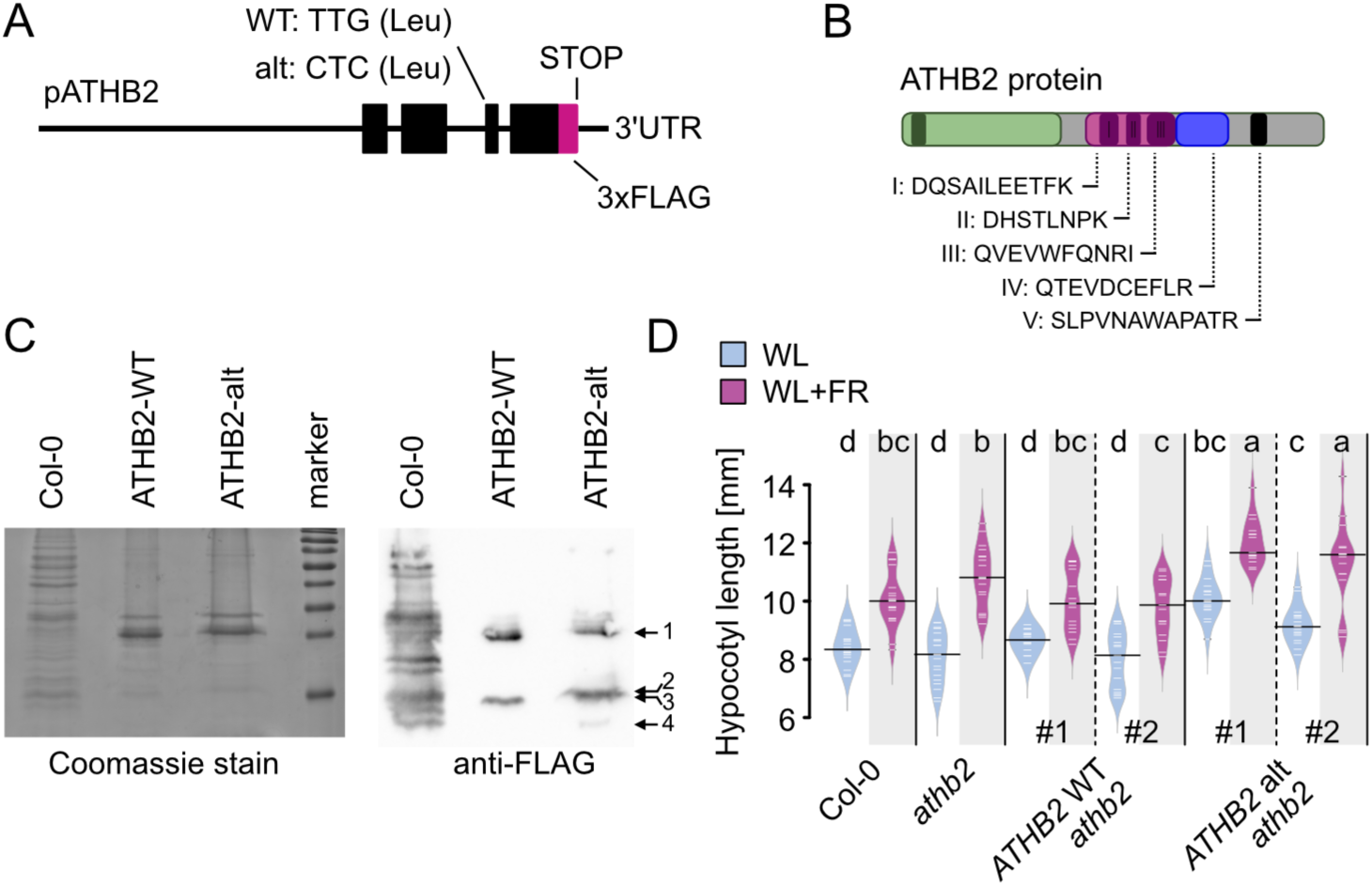
The ATHB2miP isoform restrains shade-induced elongation growth. **(A)** Schematic of the *ATHB2 WT* and *ATHB2 alt* complementation constructs introduced into the t-athb2 background. *ATHB2 alt* carries a TTG-to-CTC substitution at the alternative start codon, abolishing ATHB2miP translation without altering the full-length protein sequence. **(B)** Schematic of the ATHB2 protein depicting the positions of peptides identified by immunoprecipitation mass spectrometry. Peptides I–IV were detected exclusively in athb2 ATHB2 WT plants; peptide V exclusively in athb2 ATHB2 alt plants. **(C)** Immunoprecipitated material from Col-0, *athb2 ATHB2 WT*, and *athb2 ATHB2 alt* seedlings resolved by SDS-PAGE. Left: Coomassie-stained gel confirming equal loading. Right: anti-FLAG immunoblot. *ATHB2 WT* plants show bands corresponding to full-length ATHB2 (band 1, ∼35 kDa) and ATHB2miP (band 3, ∼15 kDa). *ATHB2 alt* plants retain full-length ATHB2 (band 1) but band 3 is replaced by two additional proteoforms of distinct sizes (bands 2 and 4), absent from *ATHB2 WT* plants. **(D)** Hypocotyl length of the indicated genotypes under white light and deep shade (13 µmol m⁻² s⁻¹; R:FR = 0.13). *ATHB2 WT* complementation restores wild-type growth; *ATHB2 alt* complementation results in exaggerated elongation under both conditions. Letters denote statistically distinct groups (Tukey HSD, p < 0.05).

To determine whether ATHB2 and ATHB2miP could be detected at the protein level, we subjected *athb2 ATHB2 WT* and *athb2 ATHB2 alt* plants to one hour of shade treatment, isolated nuclei, and performed anti-FLAG immunoprecipitation followed by mass spectrometry analysis. Several peptides corresponding to ATHB2 were identified. Notably, peptides I-IV (Fig. 5B) were detected exclusively in *athb2 ATHB2 WT* plants, while peptide V was detected exclusively in *athb2 ATHB2 alt* plants, indicating that the TTG-to-CTC substitution fundamentally alters which ATHB2 proteoforms are produced.

To corroborate these findings at the protein level, immunoprecipitated material was resolved by SDS-PAGE and analysed by immunoblotting. Coomassie staining of the gel confirmed equal loading across samples. Col-0 wild-type plants, which lack a FLAG-tagged transgene, yielded numerous non-specific bands in the immunoprecipitate, whereas *athb2 ATHB2 WT* and athb2 ATHB2 alt plants showed substantially fewer bands, consistent with efficient and specific FLAG immunoprecipitation (Fig. 5C). Immunoblotting against FLAG revealed two major bands in *athb2 ATHB2 WT* plants (labelled 1 and 3), corresponding to full-length ATHB2 (35 kDa) and ATHB2miP (15 kDa), respectively. In *athb2 ATHB2 alt* plants, full-length ATHB2 (band 1) was also detected, but the ATHB2miP band (band 3) was replaced by two distinct bands: band 2, which migrates slightly above the expected ATHB2miP position, and band 4, which migrates at approximately 5kDa, below the expected ATHB2miP size. The introduction of the TTG-to-CTC substitution therefore does not simply abolish production of the short isoform but redirects translation to alternative initiation sites, generating proteoforms of distinct sizes that are absent from *ATHB2 WT* plants.

To determine whether the loss of the TTG-initiated ATHB2miP isoform affects shade responsiveness, we analysed hypocotyl growth under white light and deep shade conditions. As observed previously, *t-athb2* mutants displayed altered hypocotyl elongation compared to wild-type plants (Fig. 5D). Complementation with the *ATHB2 WT* construct largely restored wild-type hypocotyl length under both conditions. In contrast, complementation with *ATHB2 alt* resulted in a robust and reproducible increase in hypocotyl elongation, exceeding that of wild-type, *t-athb2*, and *ATHB2 WT*-complemented lines, particularly under shade (Fig. 5D). Consistent with this, ATHB2 protein levels were slightly reduced in *athb2 ATHB2 alt* plants, suggesting that the alternative proteoforms produced in this line are less effective at relieving ATHB2 autorepression. Together, these results demonstrate that the TTG-initiated ATHB2miP isoform is required to restrain shade-induced elongation growth, supporting a model in which ATHB2miP acts as a negative regulator of ATHB2 activity.

### The shade avoidance response is influenced by nutritional status regulated by ATHB2

ATHB2 regulates the expression of genes involved in shade avoidance and root development (Fig. 3) and mutations in *ATHB2* result in defects in these processes (Fig. 4). After observing an increase in the expression of genes involved in iron transport and metabolism (Supplementary dataset 2), we investigated whether iron content was altered in *athb2* mutants and whether iron affected shade-induced hypocotyl elongation and root development.

Iron homeostasis is under the regulation of a transcriptional cascade involving genes encoding iron transporters and a family of small peptides involved in iron deficiency signalling (Spielmann *et al*, 2023). The basic helix-loop-helix family of transcription factors, specifically bHLH38, bHLH39, bHLH100, and bHLH101, have been identified as intermediaries in iron deficiency signaling for the iron transporter IRT1. They achieve this by dimerizing with FIT1, FER-LIKE IRON DEFICIENCY INDUCED TRANSCRIPTION FACTOR1 (Colangelo & Guerinot, 2004).

Transcriptome profiling of wild type, *t-athb2* and *35S::miP* seedlings revealed genes that were upregulated in both *t-athb2* and *35S::miP* plants (Fig. 3B), indicating that these genes are repressed by ATHB2. The iron deficiency cascade, which includes *bHLH38*, *bHLH39*, *bHLH100*, *bHLH101*, *FIT1*, the iron transporter *IRT1*, and the *IRONMAN* genes (*IMA1*, *IMA2*, and *IMA3*), was among the ATHB2 repressed genes and found to be upregulated in *35S::miP* and *t-abth2* seedlings (Fig. 6A,B). In white light conditions, a smaller set of genes, including *bHLH38*, *bHLH101*, *IMA1*, and *IMA2*, are already highly expressed (Fig. 6A). The remaining genes are upregulated after 45 minutes of additional far-red light (Fig. 6B), suggesting negative regulation by ATHB2 in response to shading. This regulation is uncoupled in *t-athb2* and *35::miP* plants. To determine whether the observed increases in transcription of genes encoding iron signaling and uptake components are related to iron levels, we measured the iron content of wild type, *35S::miP*, and *t-athb2* mutant seedlings. Iron is taken up by the roots and then distributed throughout the plant, and we therefore decided to measure iron levels in both roots and shoots. As anticipated, higher concentrations of iron were found in the roots than in the shoots (Fig. 6C). A two-way ANOVA revealed a strong effect of tissue, with values consistently higher in roots than in shoots across all genotypes (p < 0.001). No significant main effect of genotype and no genotype by tissue interaction were detected, indicating that the tissue difference was largely conserved among genotypes. To resolve genotype effects within tissues, we performed Tukey post-hoc comparisons. In both shoots and roots, *35S::miP* plants exhibited significantly higher iron levels than WT, whereas *t-athb2* plants showed intermediate levels that did not differ significantly from either genotype. Thus, although tissue identity is the dominant determinant of variation, the *35S::miP* genotype is associated with elevated iron levels relative to WT in both tissues. These findings collectively support the role of ATHB2 as a negative regulator of iron uptake.

**Figure 6.**
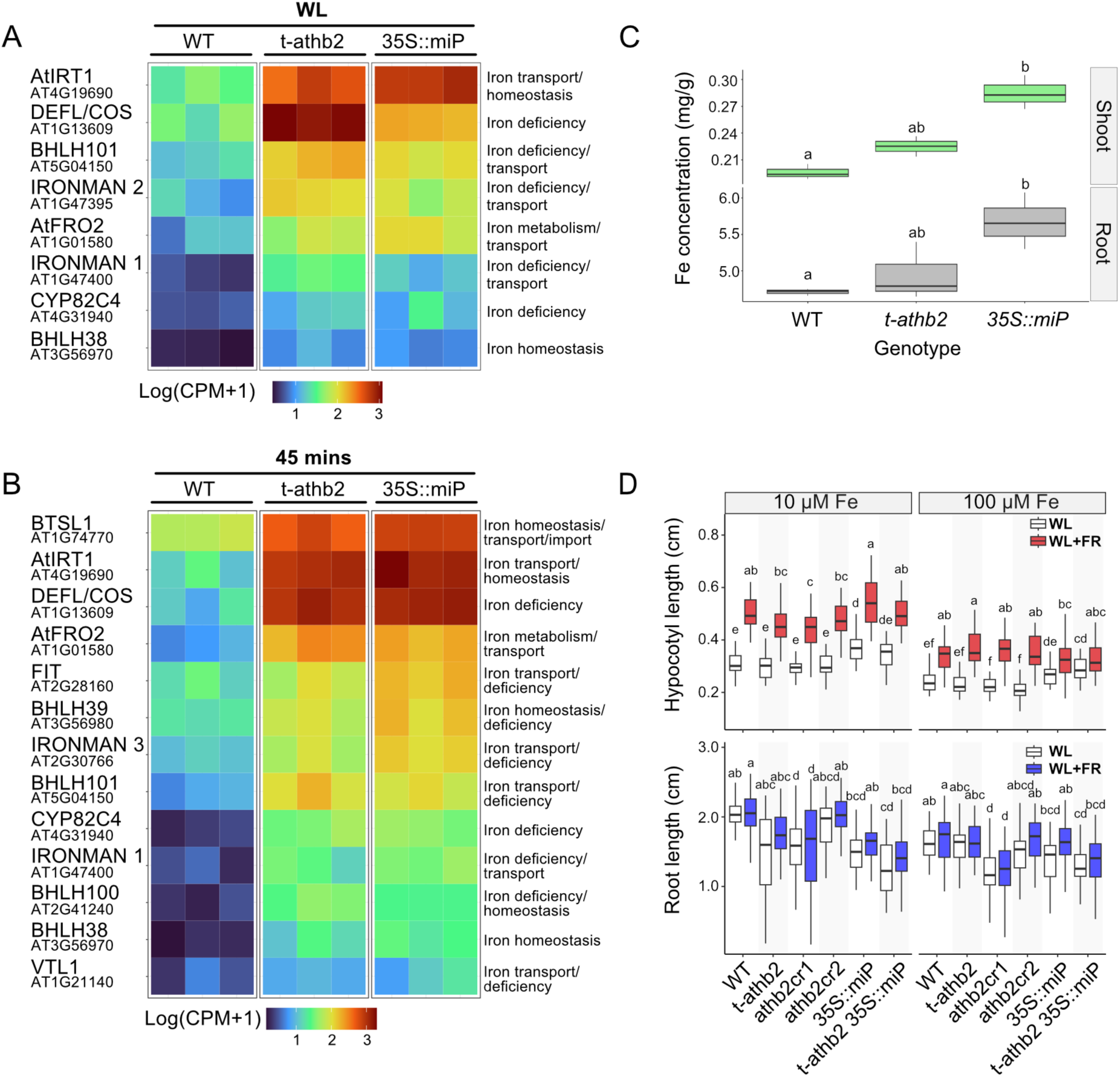
ATHB2 and ATHB2miP have a role in iron transport regulation and homeostasis. **(A)** Heatmap showing the expression levels in white light of genes upregulated in both the *t-athb2* mutant and the *35S::miP* lines that are involved in iron transport, iron homeostasis and iron deficiency response. Lowly expressed genes are coloured blue, highly expressed genes are coloured red. Additionally, the expression level for all three replicates per genotype is displayed. **(B)** Heatmap showing the gene expression levels in shade conditions. **(C)** Concentration of Fe (mg/g) measured in the roots (grey) and the shoots of seedlings of the three genotypes (green). Letters in all signify results of pairwise comparisons by two-way ANOVA. **(D)** Plots of hypocotyl length (red) and root length (blue) of plants grown on low or high iron supplemented ½ MS media in WL and WL+FR. Hypocotyls were measured after five days in either condition whilst roots were measured after nine days.

To test whether nutrient status affects root development and shade avoidance responses, we performed shade avoidance assays with seedlings grown on low iron (10mM) and high iron (100mM) media. Overall, we observed a significant hypocotyl shortening and reduced hypocotyl response when seedlings were grown on high iron media compared to low iron media. The responses to deep shade on low-iron medium were similar to those observed earlier (Fig. 4C), except that *t-athb2* mutants did not elongate hypocotyls under white light conditions. On high iron media, we observed a dampening of the shade avoidance response, but *t-athb2*, *athb2cr2* and *athb2cr1* mutants showed longer hypocotyls than wild type under shade conditions. Transgenic plants ectopically expressing ATHB2miP (35S::miP and *t-athb2 35S::miP*) showed slightly elongated hypocotyls in white light and almost no elongation in shade. These results show that nutrient status has a strong influence on the shade avoidance response and that the function of ATHB2 is dependent on iron homeostasis. Under low iron conditions, ATHB2 acts as an activator of shade growth, whereas under high iron conditions it suppresses shade growth. With regard to root growth, we measured the length of the primary root on low and high iron media in the absence and presence of shade. Overall, we did not observe strong changes in root elongation between white light and shade grown seedlings, supporting what we had observed previously (Fig. S4). However, we observed that all *athb2* mutant plants, with the exception of *athb2cr2* mutants, developed significantly shorter roots when grown on low-iron medium. This finding may indicate that these mutants, in which iron uptake is strongly up-regulated, do not need to grow longer roots to increase their iron uptake. The latter hypothesis is supported by the finding that plants ectopically expressing ATHB2miP have the shortest roots but show the greatest hypocotyl elongation in shade. On high-iron medium, we did not observe major changes in root length, except for *athb2cr2* mutants, which had significantly shorter roots in both white light and shade conditions. In conclusion, these results show that nutrient status affects both root length and hypocotyl elongation in response to shade. ATHB2 mediates between root growth and hypocotyl elongation and can either activate or repress hypocotyl elongation in response to different iron concentrations.

## Discussion

In this study, we assessed the impact of shading on transcript isoform production and their potential to encode microproteins. It was found that almost half of the Arabidopsis genes that had reads at the transcription start site also had alternative transcription start sites in the gene body. Only a fraction of these genes had the potential to encode microproteins. It is furthermore fair to assume that many more potential microproteins exist, but these are produced in a tissue-specific and/or inducible fashion. We filtered the dataset to prioritize alternative transcripts with robust expression levels, excluding those unsupported by multiple reads, which again could be candidates that have a narrower pattern of expression. Our approach focused on enriching microprotein candidates likely to have biological significance. To enhance the likelihood of identifying microproteins involved in the shade avoidance response, we considered alternative methods for studying transcript isoforms, including parallel analysis of RNA 5′ends from low-input RNA (nanoPARE), and ribosome profiling (Ingolia, 2014; Schon *et al*, 2018). Ribosome profiling, which employs deep sequencing to monitor *in vivo* translation, can provide additional insights into protein formation. Such future approaches can both confirm the purely sequencing-based identification of candidate microproteins but also aid to identify novel candidates.

The list of microprotein candidates generated through the 5’PEAT-seq approach contained few shade avoidance regulators (Supplementary dataset 1). Notably, microproteins may serve distinct functions compared to their full-length counterparts, albeit often targeting evolutionarily related proteins. Adjusting filtering criteria could identify more transcript isoforms related to shade responses, although they may not classify as microproteins. The ATHB2miP microprotein, chosen due to its similarity to the known LITTLE ZIPPER microproteins (Kim *et al*., 2008; Wenkel *et al*., 2007), was expected to interact with ATHB2 and regulate its activity, akin to ZPRs and HD-ZIPIIIs. Yeast-Two-Hybrid tests and pull-down assays confirmed physical interactions between ATHB2 and ATHB2miP. Electrophoretic mobility shift assays further demonstrated that, similar to LITTLE ZIPPER microproteins (Wenkel *et al*., 2007), ATHB2miP interferes with the DNA-binding activity of the full-length ATHB2 protein by sequestering it into a non-DNA-binding complex. Notably, ATHB2miP can also homodimerize, a feature attributed to the leucine zipper domain’s role as a protein-protein interaction motif in HD-ZIP proteins.

We assessed potential alternative *ATHB2miP* promoters using the GUS protein assay, identifying one active promoter, *pATHB2miP::GUS*, which did not possess a typical ATG start codon. This observation is consistent with prior findings of alternative start sites not commencing with an ATG. Differences in GUS expression patterns between *ATHB2* and *ATHB2miP* promoters were observed, with ATHB2miP primarily present in the shoot apical meristem (SAM) and early leaf primordia. The absence of ATHB2miP in other tissues suggested that it does not directly regulate ATHB2 in those areas.

Analysis of the subcellular localization patterns of ATHB2miP and ATHB2 protein-fusions to GFP revealed that ATHB2 localizes mostly to larger subnuclear speckles, a phenomenon also observed for phytochromes and other light signaling components and therefore often referred to as photobodies (Kim *et al*, 2023; Kircher *et al*, 2002). Interestingly, ATHB2miP protein showed a more diffuse nuclear localization implying discrete functions of both isoforms. However, when co-expressed both isoforms appear in speckles, but these are more dispersed, indicating a direct physical interaction where the full-length isoform is partially removed from the speckles. The full-length ATBH2 isoform has an intrinsically disordered amino terminus and could undergo phase separation and engage in more flexible protein-protein interactions within the photobodies. As ATHB2miP lacks the disordered part, it may prevent the phase separation activity of ATHB2.

To investigate the regulatory role of ATHB2miP, we compared *ATHB2miP* overexpression plants to *athb2* mutants. Surprisingly, in deep shade conditions all *athb2* knock-down and knock-out mutants as well as *ATHB2miP* overexpression plants showed elongated hypocotyls in white light conditions compared to Col-0 wild type plants while the hypocotyl length in shade grown seedlings showed no significant differences when compared to wild type. Except 35S::miP transgenic plants that showed excessive hypocotyl elongation in both white light and far-red enriched conditions (Fig. 4C). This indicates that ATHB2 functions as a growth repressor in deep shade and ATHB2miP has the capacity to inactivate ATHB2 which results in hypocotyl elongation in shade. The complementation analysis provided direct genetic evidence for a regulatory role of ATHB2miP in controlling elongation growth. Preventing ATHB2miP translation by mutating the alternative start codon resulted in exaggerated hypocotyl elongation, both in white light and under deep shade, whereas complementation with the wild-type *ATHB2* locus restored growth responses to near wild-type levels (Fig. 5D).

The proteomics and western blot data from the complementation lines reveal an unexpected complexity in ATHB2 proteoform production. When the TTG start codon is mutated, translation does not simply cease but shifts to alternative initiation sites, producing proteoforms of different sizes that are absent from *ATHB2 WT* plants. This suggests that the *ATHB2 l*ocus has an intrinsic capacity to generate multiple short isoforms, with the TTG codon normally acting as the preferred and dominant initiation site, likely due to its favorable Kozak context. The alternative proteoforms produced in *ATHB2 alt* plants are evidently less effective at restraining ATHB2 activity, as reflected in the exaggerated elongation phenotype of these lines. Whether this reflects altered leucine zipper conformation, reduced interaction efficiency with ATHB2, or impaired access to photobodies remains to be determined. The non-specific bands observed in Col-0 immunoprecipitates are consistent with the absence of a FLAG-tagged bait protein, which renders the immunoprecipitation non-selective; the substantially cleaner profiles in the complemented lines confirm that the identified bands represent genuine FLAG-tagged ATHB2 proteoforms rather than background.

Given the transient induction of the *ATHB2miP* transcript upon shade exposure (Fig. 1B,C), these results support a model in which ATHB2miP acts as an early inhibitory brake on ATHB2 activity, restraining premature or excessive growth during short-term shading. As shade persists, ATHB2miP expression declins, thereby relieving this inhibition and allowing sustained elongation responses. Abolishing this regulatory constraint consequently resulted in excessive growth in both white light and shade conditions. The finding that the growth differences can only be observed in deep shade conditions indicates that ATHB2 activity is dependent on other components that are only present in this type of shade. In canopy and proximity shade ATHB2 functions as a promoter of elongation growth, hence the shorter hypocotyls in these types of shade.

ATHB2 has also been reported to be involved in controlling root growth and development (Schena *et al*., 1993) prompting us to investigate the role of ATHB2 and ATHB2miP in this process. The analysis of root development in different types of shade did not reveal significant differences in root length. However, there were notable differences in the ability of *athb2* mutants to produce lateral roots. Additionally, all tested *athb2* mutants appeared to be insensitive to shade, as the number of lateral roots did not change in response to proximity shade, unlike in the wild type. In summary, these results suggest that ATHB2 acts as a suppressor of lateral root growth in response to proximity shade. The overexpression of ATHB2miP had a significant impact on lateral root development in both white light and shade. Notably, in the *t-athb2* mutant background (*t-athb2 35S::miP*), the strong shade suppression was not observed, indicating that ATHB2 is required for strong shade suppression. These findings suggest that ATHB2miP has a role beyond ATHB2 inhibition. The discovery that ATHB2 has diverging functions depending on the type of shade experienced by the plant raises questions about the upstream signalling processes that control ATHB2 activity and the type of protein complexes in which ATHB2 participates. Additionally, the localization of ATHB2 to photobodies hints at a role in phytochrome B signalling. In deep shade, phyA may have additional roles that cause ATHB2 to switch from being a growth activator to being a growth inhibitor. To better understand ATHB2’s molecular functions in different light conditions, it is important to study the protein complexes it interacts with.

To elucidate the molecular events resulting from the absence of *ATHB2* or the inhibition of ATHB2 by overexpression of ATHB2miP, we conducted an RNA-sequencing analysis comparing Col-0 wild type with *t-athb2* and transgenic plants overexpressing ATHB2miP (*35S::miP*), all grown under both white light and deep shade conditions. In line with the exaggerated growth phenotype of the hypocotyl in deep shade, we found auxin-and shade-related genes to be expressed at elevated levels. The fact that the expression of these genes follows the same trend in both the *t-athb-2* and the *35S::miP* line also supports the negative feedback mechanism we hypothesised. In addition, we also found genes involved in root growth and development to be over-represented in white light grown seedlings, which corresponds to what we observe in our lateral root phenotyping results. Genes involved in the formation of lateral roots were found to be upregulated in both the *t-athb2* and the *35S::miP* seedlings, even in shade. Using gene clustering analysis, especially cluster 2 (Fig. 3B) comprises genes that are down-regulated in both the *t-athb2* and the *35S::miP* lines and several of these genes are involved in root growth and development. Surprisingly, genes that were upregulated in *t-athb2* and *ATHB2miP* overexpression plants encode iron transporters. The finding that iron concentrations were elevated in these plants implies that these roots might have stopped growing because of a better nutrient supply. Our findings show that iron availability influences the shade avoidance response and primary root length. Furthermore, it appears that the nutritional status influences the activity of ATHB2, suggesting that ATHB2 is integrating different environmental inputs (shade and nutrient availability) to formulate an appropriate response that can either activate or repress growth.

Shade not only promotes elongation growth, but it also affects the plant’s hormone homeostasis, weakening its ability to defend against pathogens (de Wit *et al*, 2013). The recent discovery that regulation of iron uptake is influenced by internal IRONMAN peptides, which are sensitive to external flagellin perception (Cao *et al*, 2024), raises the question of whether ATHB2 may play a central role in mediating between nutritional immunity and shade-induced growth promotion or suppression.

In summary, our work uncovered transcript isoforms that are responsive to shading, including ATHB2miP, a microprotein derived from an alternative transcription start site in the *ATHB2* gene. This microprotein appears to intercede between root and shoot growth in response to shade. In summary, our data support a model for ATHB2/ATHB2miP as integrator of nutrient status and elongation growth of both the shoot and the root.

## Materials and methods

### Plant materials and growth conditions

All plants were grown in growth chambers on ½ Murashige and Skooge (MS) 1% agar under long day (LD) conditions at 22°C during daytime and 20°C during night. For the shade treatment for the 5’PEAT-seq and RNA-seq experiments, seedlings were transferred after 10 days from a white light only compartment (PAR of 13 μmol/m^2^/s, R:FR = 7.7) to a compartment with additional far-red lights (PAR of 13 μmol/m^2^/s, R:FR = 0.2) for treatments of 0, 45 and 90 minutes. Whole seedlings frozen in liquid N_2_ before grinding with a mortar and pestle and RNA extraction. For the shade treatments related to measuring hypocotyl length and root growth, half of the plants were transferred after two days from a white light only compartment (PAR of 45 μmol/m^2^/s, R:FR = 5.5) to a compartment in the same chamber with additional far-red lights (PAR of 45 μmol/m^2^/s, R:FR = 0.15). The other half remained in the white light only compartment. After five and eleven days, seedlings were photographed and hypocotyls and roots measured using IMAGEJ.

For the growth and phenotyping of seedlings on iron-enriched and iron-depleted media, two different growth media were prepared. Both media contained the same vitamins, macro and micro elements of standard MS salt (M0222, Duchefa) and in the same concentrations, with the exception of FeNAEDTA, used in a concentration of 10uM in the iron-depleted medium and 100uM in the iron-enriched medium. Seedlings were grown on 1% agar iron-depleted and iron-enriched media under the same growth conditions of the other growth experiments. Half of the seedlings on plates were transferred after two days from a white light only compartment (PAR of 13 μmol/m2 /s, R:FR = 7.7) to a compartment in the same chamber with additional far-red lights (PAR of 13 μmol/m2 /s, R:FR = 0.2). The other half remained in the white light only compartment. After five and eleven days, seedlings were photographed and hypocotyls and roots measured using IMAGEJ.

### 5’-PEAT library preparation and analysis

Arabidopsis Col-wild type seedlings were grown for 10 days on 1/2 MS agar plates as described before. Whole seedlings were harvested directly into liquid nitrogen after 0, 45 or 90 minutes of shade treatment. Total RNA was extracted with Spectrum Plant Total RNA Kit (Sigma). PEAT library preparation was done exactly as described earlier (Ni *et al*., 2010), but all steps beginning with the circularization of the PCR product were omitted and the fragments were inserted into NEBNext Ultra™ II DNA Library Prep K it for Illumina and quantified with NEBNext Library Quant Kit for Illumina. The finished libraries had a fragment size of 500 bp. BGI tech solutions (Hongkong) performed sequencing on an Illumina HiSeq (150bp paired end, 40 M read pairs per sample).

Quality control was done with FastQC (v0.11.5), available online at: http://www.bioinformatics.babraham.ac.uk/projects/fastqc) and generic Illumina adapters were removed with Cutadapt (Martin, 2011). Additionally, Cutadapt removed PEAT-adapters and marked read direction for later identification of the RNA start. HISAT2 (v2.0.5) (Kim et al., 2015) aligned the reads with more than 90% success to the Arabidopsis thaliana TAIR9 genome assembly (Lamesch *et al*, 2012). Samtools (v0.1.19) (Li *et al*, 2009) and bedtools (v2.17.0) (Quinlan & Hall, 2010) filtered for properly paired reads with high aligment score (-q 10) and performed file conversions. Around 30 million read pairs per sample were further analysed. Using previously marked read directions, RNA starts were identified and recorded in bedgraph format. Only genes with a transcription start site (TSS) in the 5’UTR or upstream region and at least one alternative TSS in coding sequence or intron region were further analysed. The difference between the TSS and the alternative start site was set to 10%. To test whether the regulation of the 5’UTR-TSS and the alternative TSSs are different, the number of TSS-reads in the 5’UTR-TSS and one alternative TSS were compared in two samples with Fisher’s exact test and Benjamini-Hochberg corrected false discovery rate in R software (v3.2.2). The protein size was set between 7 and 20 kDa. Pfam-A domains (v28, ftp://ftp.ebi.ac.uk/pub/databases/Pfam/) (Finn *et al*, 2016) were searched in protein sequences with hmmer3 (http://hmmer.org/) and cutoffs of e-value <= 0.1 and c-value <= 0.05. Interaction domains were identified using iPfam (v1.0, http://www.ipfam.org/) (Finn *et al*, 2014).

### Generation of transgenic and CRISPR-Cas9 edited plants

To generate the overexpression of ATHB2miP (*35S::ATHB2miP*) in Arabidopsis oligos were designed to amplify exons 3 and 4 of *ATHB2* and the resulting sequence then cloned into the vector pJAN33 by gateway cloning. Plants were transformed by floral dipping and transformants selected for by BASTA treatment.

For CRISPR mutagenesis, sgRNAS were cloned into the CRISPR/Cas9 vector pKI1.1R. The cloning was performed as described (Tsutsui and Higashiyama, 2016). *Athb2cr2* was made by Cas9 gene editing targeted using the following sgRNA combination: 5’-TACAGTCTCAAGCTCTACA-3’ and 5’-GTTTCAGAACAGACGAGCA-3’. *Athb2-cr3* was made by Cas9 gene editing targeted using the single sgRNA: 5’-ACGATCTGGGTCTAAGCTT-3’. Plants were transformed by floral dipping and then RFP-positive seeds selected for in the T1 generation. Genomic DNA of 4-week old plants was isolated using the Edwards method and the plants were genotyped by PCR. The PCR products were gel purified using the E.Z.N.A ® Cycle-Pure Kit (Omega Bio-tek) and sequenced at Macrogen (Amsterdam). Expression in transgenic and mutant plants was measured by RT-qPCR. First, RNA was extracted from 4-week old plants using the Spectrum Plant Total RNA Kit (Sigma). Purified RNA (1 μg) was used for reverse transcription using the iScript™ cDNA Synthesis Kit (BIO-RAD). RT-qPCR was performed using SYBR green (ThermoScientific) on a Biorad CFX384, using oligos against *ATHB2*: Region A FWD 5’-GATTGCAAAATCTCTCTCTCTCTC-3’ with REV 5’-GACCCAGATCGTCTTTCTCG-3’; Region B FWD 5’-TGTTCGAGAAAGACGATCTGG-3’ with REV 5’-GTCGATTCCTCGGATGAAAG-3’; Region C FWD 5’-CAGTGACGATGAAGATGGTGA-3’ with REV 5’-GAAACCAAACTTCCACTTGTCT-3’; Region D FWD 5’-GCTGAAGCAAACGGAGGTAG-3’ with REV 5’-GGACCTAGGACGAAGAGCGT-3’. Gene expression levels were standardized to the housekeeping gene GAPDH (FWD 5’-AAAGTGTTGCCATCCCTCAA-3’ with REV 5’-TCGGTAGACACAACATCATCCT-3’) using the Pfaffl equation.

### Yeast-2-Hybrid

For the Y2H assay, the coding sequences of *ATHB2* and *ATHB2miP* (exons 3 and 4 of *ATHB2,* as above) were recombined into vectors pGBKT7-GW and pGADT7-GW by gateway cloning. The constructs were transformed to yeast strains PJ694α and YM4271A using lithium acetate and heat shock for 15 min (42°C). Yeast colonies that grew after two days at 30°C on SD dropout medium plates lacking either tryptophan (-W) or leucine (-L) were selected for, mixed with each other and spread on SD dropout medium lacking tryptophan and leucine (-LW). Colonies were finally plated on dropout-LW medium that additionally lacked histidine (–LWH), with additional 10 mM 3-aminotriazole to test for protein interactions.

### Electromobility shift assay (EMSA)

DNA–protein interactions were analyzed using the Gelshift Chemiluminescent EMSA Kit (Active Motif) according to the manufacturer’s instructions. Double-stranded oligonucleotides containing the ATHB2-binding motif (CAATGATTG) were synthesized and biotin-labeled. Binding reactions (20 µL) contained 0.44 µM biotin-labeled probe, 0.04 mg/mL recombinant GST-ATHB2 protein, and increasing concentrations of recombinant GST-ATHB2-miP (0.01–0.16 mg/mL). Reactions were incubated for 20 min at room temperature. Complexes were resolved on a native DNA retardation gel, transferred to a nylon membrane, and detected using streptavidin–HRP with chemiluminescent substrate. Signals were visualized using an Azure c600 Imaging System (Azure Biosystems).

### Protein extraction and western blot analysis

Transgenic *athb2 ATHB2 WT* and *athb2 ATHB2 alt* seedlings were subjected to 60 minutes of shade treatment under the conditions used for the 5’PEAT-seq experiments described above. Approximately 1 g of seedlings was harvested and used for nuclear extraction following the method of Zhao et al. Nuclear proteins were extracted by incubating the isolated nuclear fraction in extraction buffer (20 mM HEPES pH 7.4, 300 mM NaCl, 1.5 mM MgCl2, 10% glycerol, 0.1% Triton X-100, 0.1% Nonidet P-40, 1 mM DTT, 0.5 mM PMSF, 50 U/mL DNase I, and 1x protease inhibitor cocktail) for 30 minutes at 4°C with gentle agitation, followed by centrifugation at 12,000 x g for 10 minutes at 4°C. The supernatant containing the soluble nuclear protein fraction was retained for further analysis.

Protein concentration was determined by Bradford assay. Forty micrograms of protein per sample were denatured in Laemmli sample buffer at 95°C for 5 minutes and resolved on a 12% SDS-PAGE gel alongside a pre-stained molecular weight marker. Proteins were transferred to a PVDF membrane at 70 V for 1 hour using a wet transfer system. The membrane was blocked in 5% BSA in TBST for 24 hours at 4°C, then incubated with anti-FLAG M2 antibody (Sigma; 1:4000 in 5% BSA) for 10 hours at 4°C. After three washes in TBST, the membrane was incubated with an HRP-conjugated secondary antibody (1:2000) for 1 hour at room temperature. Bands were detected using enhanced chemiluminescence and imaged on an Azure Biosystems c600.

### LC-MS/MS

Samples were digested using a modified SP3 protocol (Hughes *et al*, 2019). Briefly, samples were denatured, bound to SpeedBeads, digested overnight with trypsin, and desalted. Peptides were separated on a Vanquish Neo (Thermo) system equipped with a PEPMAP NEO C18 trapping column (300 µm × 5 mm, 5 µm) and a nanoEase M/Z HSS C18 T3 analytical column (75 µm × 250 mm, 1.8 µm) and introduced into an Exploris 480 mass spectrometer (Thermo). Data were acquired in data-dependent mode with survey scans covering m/z 375–1500 at 120,000 resolution, RF lens 40%, and normalized AGC of 300%. A maximum cycle time of 2 s was used to select precursors of charge states 2–6 for MS2 analysis. Dynamic exclusion was set to 35 s. MS2 scans were acquired at 15,000 resolution with auto AGC, 1.4 m/z isolation window, and HCD fragmentation at NCE 30. Isotopes were excluded from MS2 selection.

Raw data were searched against the Arabidopsis thaliana UniProt proteome (UP000006548) appended with ATHB2 isoform sequences using FragPipe (v18) with the LFQ-MBR workflow for label-free quantification. Contaminant and decoy hits were removed prior to analysis.

### GUS localization assay

Oligos were designed to amplify the 1 kb sequence upstream of *ATHB2,* and the sequences upstream of the alternative start codon TTG in exon 3 and upstream of a control start codon ATG in intron 2, up to and including the putative start codon ATG of *ATHB2.* Sequences were cloned into pBGWFS7 by gateway cloning and the resulting vectors transformed into plants by floral dipping. Arabidopsis thaliana (ecotype Col-0) seeds were surface-sterilized and sown on Murashige and Skoog (MS) agar plates. The plates were incubated at 4°C for 2 days for stratification and subsequently transferred to a growth chamber under long-day conditions. Seven day old seedlings were immersed in GUS staining solution (XGLUC; 14-mM NaH2PO4, 36-mM Na2HPO4, 2-mM K4[Fe(CN)6], 2-M K3[Fe(CN)6], 10% Triton X-100). Seedlings were vacuum infiltrated for 5 minutes, incubated overnight at 37 °C in the dark, and rinsed with PBS to remove excess staining solution and dehydrated through an ethanol series (30%, 50%, 70%, 90%, and 100%) to remove chlorophyll and enhance contrast.. Pictures of seedlings were taken with a canon EOS 450D camera on a Nikon Eclipse 80i Fluorescence microscope. White balance and scalebar were added via ImageJ.

### Subcellular localization and FRET-FLIM analysis

As above, the coding sequences of *ATHB2, ATHB2miP* and *ATHB2HD* (from exon 1 to the beginning of exon 4) were cloned into the vectors pK7WGF2 that contains an N-terminal eGFP driven by a CaMV35S promoter and a modified vector of pEarlyGate104 containing an N-terminal mCherry driven by a CaMV35S promoter (provided by Sabine Müller/Dorothee Stöckle, ZMBP Tübingen). Tobacco leaves were transiently transformed by agrobacterium infiltration (three individual plants per infiltration) and nuclei of the epidermal cells of leaves were imaged 2-3 days post-infiltration using a Leica Stellaris 8 confocal laser-scanning microscope. All vectors were co-transformed with p19 and for the FRET-FLIM assay, eGFP-ATHB2 was additionally co-transformed with either mCherry, mCherry-ATHB2, or mCherry-ATHB2miP. Fluorescence lifetime measurements were collected using the TauInteraction tool in the LAS X software; samples were excited with a 470 nm pulsed laser (10 MHz) and emission from 500 nm to 560 nm was recorded.

### Differential Gene Expression Analysis and Clustering

Total RNA was extracted from Arabidopsis thaliana Col-0 wild-type (WT), t-athb2, and 35S::ATHB2miP genotypes under three different light conditions as described: White Light (WL), 45 minutes of Far-Red (FR) enrichment, and 90 minutes of FR enrichment. Three biological replicates per genotype-condition combination were used. RNA quality and concentration were assessed using a NanoDrop spectrophotometer and Agilent Bioanalyzer. RNA was sequenced on an Illumina HiSeq platform.

Raw sequencing reads were quality-checked using FastQC to ensure data integrity. Trimming of reads to remove adapter sequences and low-quality bases was performed using Trimmomatic. After preprocessing, high-quality reads were retained for downstream analysis. Read alignment to the Arabidopsis reference genome (TAIR10) was conducted using the STAR aligner within the R environment. This generated BAM files for each sample, which were subsequently used for quantification of gene expression. Differential gene expression analysis was carried out using the edgeR package in R. Raw counts were normalized using the TMM (Trimmed Mean of M-values) method. Genes with low expression were filtered out, and statistical analysis was performed to identify differentially expressed genes (DEGs) between genotypes and conditions. P-values were adjusted for multiple testing using the false discovery rate (FDR) method. DEGs were subjected to gene ontology (GO) enrichment analysis to gain insights into biological processes and pathways associated with the observed expression changes. The topGO package in R was used for this purpose.

The raw counts obtained from the RNA-seq analysis were used as input for the MFuzz clustering. These count data represented the expression levels of genes across the three genotypes and three light conditions.

For the clustering, the count data were first normalized (TMM method) to account for differences in sequencing depth and library size. MFuzz, an R package specifically designed for model-based clustering of high-dimensional gene expression data, was utilized for clustering analysis. The resulting 17 gene clusters were visually inspected to find the ones with the wished expression patterns across the samples. Genes belongind to clusters of interest were extracted and subjected to gene ontology (GO) enrichment analysis.

The reads were visualised on IGV (Integrative Genome Viewer) from the BAM mapping files generated.

### Preparation of samples and iron concentration measurements

WT, t-athb2 and 35S::ATHB2miP seedlings (about 200 per genotype) were grown on standard MS medium for 12 days. They were carefully removed from the medium to avoid damage and the aerial part of each seedling was excised from the roots. In all steps, no iron nor glass tools were used. The roots and shoots were rinsed in a 2 mM CaCl_2_ solution to remove iron accumulated on the surface. The material was placed in an Eppendorf tube and dried in a stove at 70°C for 3 days. Each of the dry samples was weighed (dry root weights ranged from 2 to 6 mg, seedlings from 15 to 25 mg). Samples were digested in 2.5 mL of 70% HNO3 + 1 mL of H2O2. Digestion conditions: maximum temperature = 240°C; maximum pressure = 200 bar; cycle duration: 1-2 hours. Following digestion, the samples were diluted in ultrapure Milli-Q water to achieve a [HNO3] = 3.5%. The liquid samples were then analyzed using an ICP-QQQ-MS 8800 Agilent instrument. NIST certified reference materials were employed to verify the accuracy of the data.

## Acknowledgements

We thank Jaume Martinez-Garcia for comments on the manuscript.

## Additional information

### Funding

We acknowledge funding through NovoCrops Centre (Novo Nordisk Foundation project number 2019OC53580 to S. W.), the Independent Research Fund Denmark (0136-00015B and 0135-00014B to S. W.) and the Novo Nordisk Foundation (NNF18OC0034226 and NNF20OC0061440 to S. W.). This work was partially supported by the Wallenberg Initiatives in Forest Research (WIFORCE) funded by the Knut and Alice Wallenberg Foundation.

## Author contribution

### Conceptualization

Stephan Wenkel.

### Data curation

Daniel Straub, Maurizio J. Chiurazzi.

### Formal analysis

Ashleigh Edwards, Anne van Humbeeck, Maurizio Junior Chiurazzi, Anko Blaakmeer, Ylenia Vittozzi, Ashish Sharma, Krishna Kodappulli Das, Sanne Matton, Valdeko Kruusvee, Daniel Straub, Giovanna Sessa, Monica Carabelli, Giorgio Morelli, Stephan Wenkel

### Funding acquisition

Stephan Wenkel.

### Investigation

Ashleigh Edwards, Maurizio Junior Chiurazzi, Anko Blaakmeer, Anne van Humbeeck, Ylenia Vittozzi, Ashish Sharma, Krishna Kodappulli Das, Sanne Matton, Valdeko Kruusvee, Daniel Straub, Giovanna Sessa, Monica Carabelli <u>Methodology:</u> Daniel Straub, Maurizio Junior Chiurazzi, Shaochun Zhu, Andre Mateus, Valdeko Kruusvee

### Supervision

Stephan Wenkel.

### Validation

Daniel Straub, Valdeko Kruusvee, Giovanna Sessa, Monica Carabelli, Giorgio Morelli

### Writing – original draft

Ashleigh Edwards, Maurizio Junior Chiurazzi, Anko Blaakmeer

### Writing – review & editing

Giovanna Sessa, Monica Carabelli, Giorgio Morelli, Stephan Wenkel.

## Additional files

**Supplementary dataset 1.** Alternative transcripts encoding potential microproteins discovered by 5’PEAT seq.

**Supplementary dataset 2.** Differentially expressed genes in wild type, t-athb2 and 35S::miP seedlings exposed to 45 and 90 minutes of shade.

**Supplementary dataset 3.** Gene clusters identified by RNA-seq.

## Data availability

All sequencing data have been deposited in NCBI’s Gene Expression Omnibus under GEO Series accession no. GSE241976, comprising RNA-seq (GSE241974) and 5’PEAT-seq (GSE241975), iso-seq (GSE267983).

## Supplementary figures

**Figure S1.**
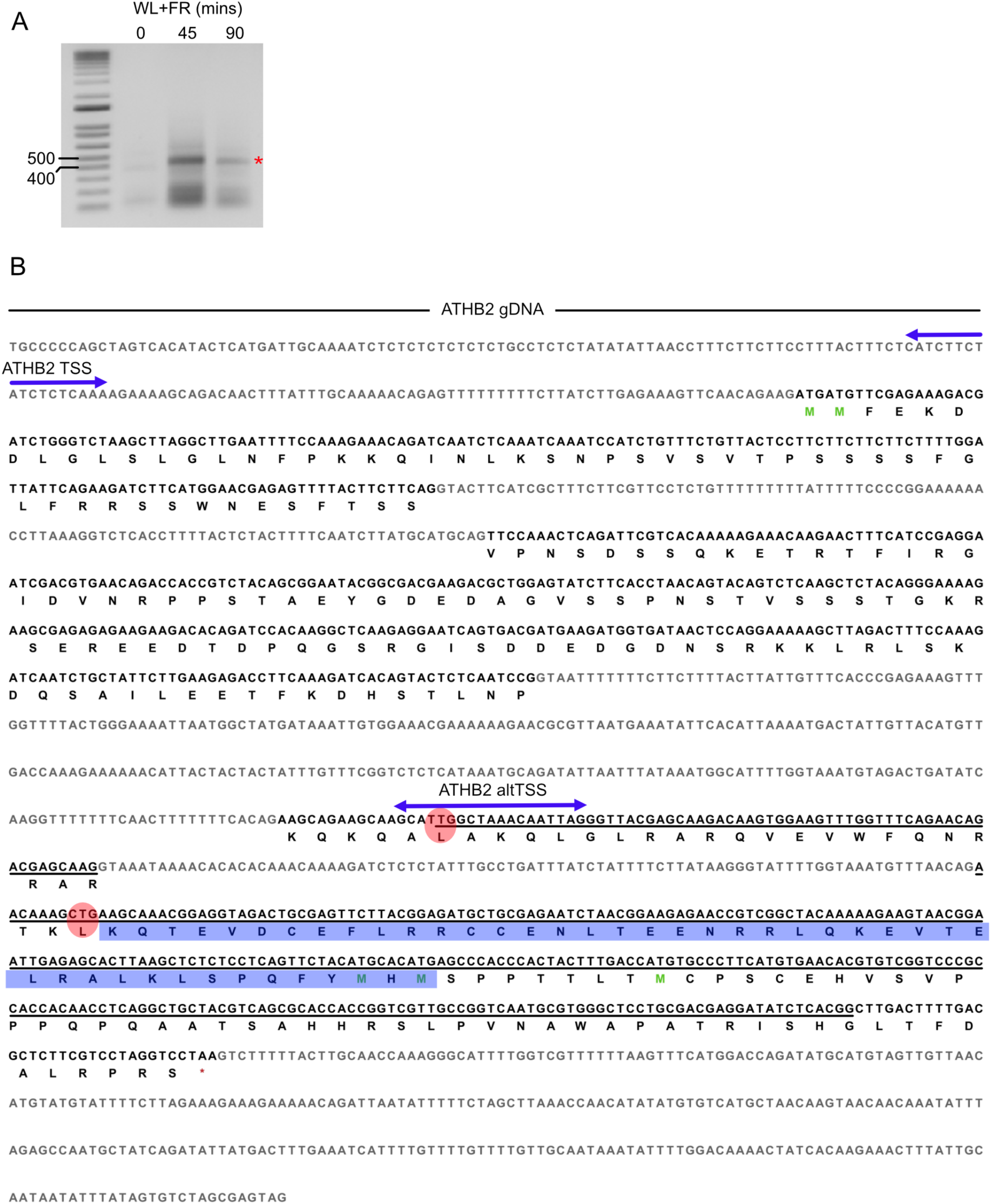
Identification of an alternative ORF in the coding sequence of *ATHB2*. **(A)** 5’RACE carried out on 10 day old seedlings that were exposed to 0, 45 and 90 mins of WL+FR light. Red asterisk signifies the small transcript arising from *ATHB2* that when purified and sequenced corresponds to the underlined portion of *ATHB2* in part B. **(B)** Genomic sequence of *ATHB2*. The exonic sequence with protein sequence below is shown in bold. Double-sided blue arrows denote sites corresponding to 5’PEAT-seq read clusters. Blue highlighted sequence denotes the leucine zipper domain. Red circles indicate two alternative non-canonical start codons identified by TIS predictor. Underlined sequence corresponds to the transcript purified from 5’RACE in part A.

**Figure S2.**
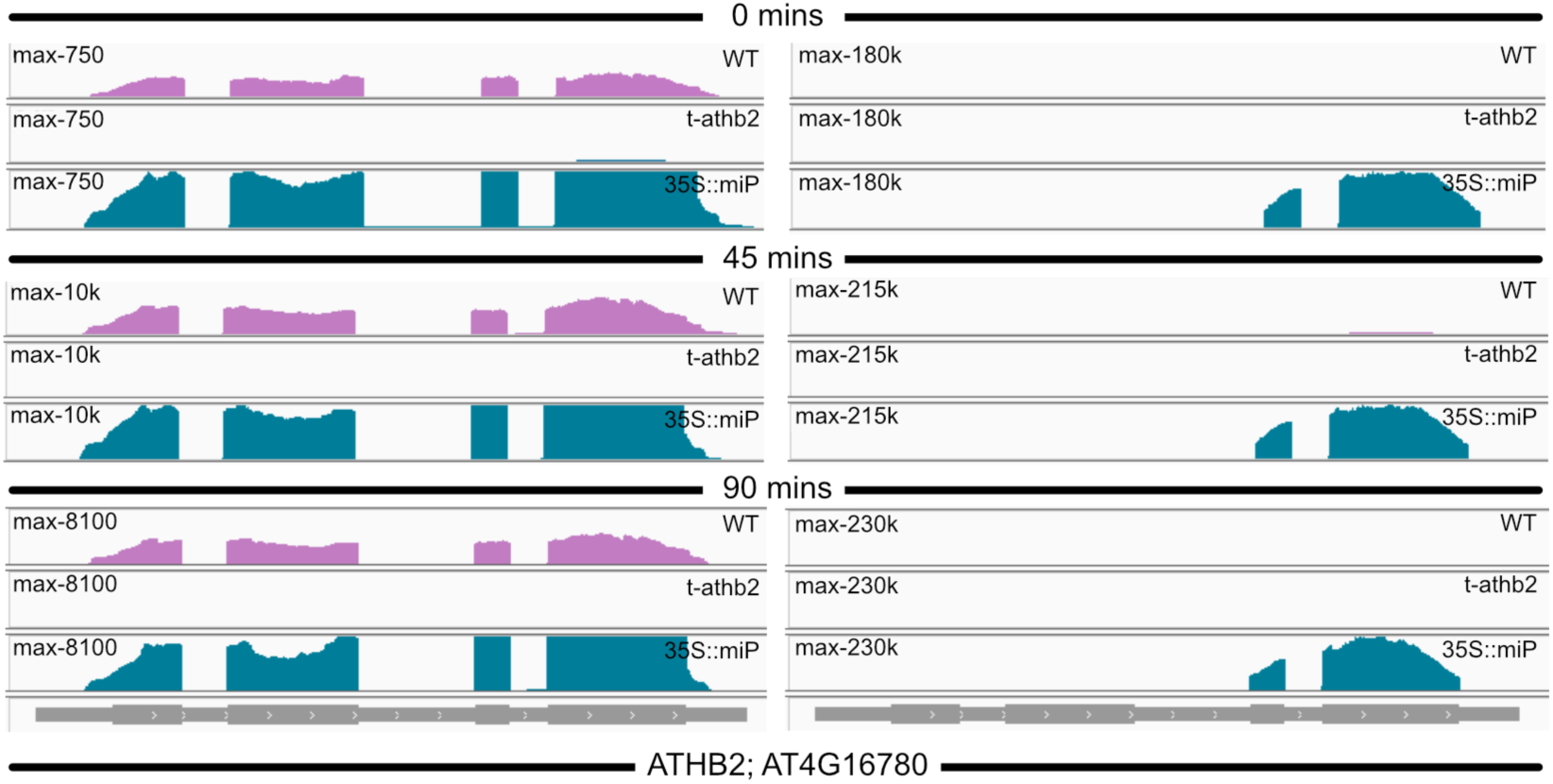
RNA-seq reads corresponding to ATHB2 at 0, 45 and 90 mins of WL+FR. RNA-seq reads peak in the WT at 45 mins of WL+FR and then decrease (see y axis for maximum reads) at 90 mins. No reads are detected in t-athb2 mutants whereas 35S::miP mutant plants are highly overexpressing exon 3 and 4 of ATHB2.

**Figure S3.**
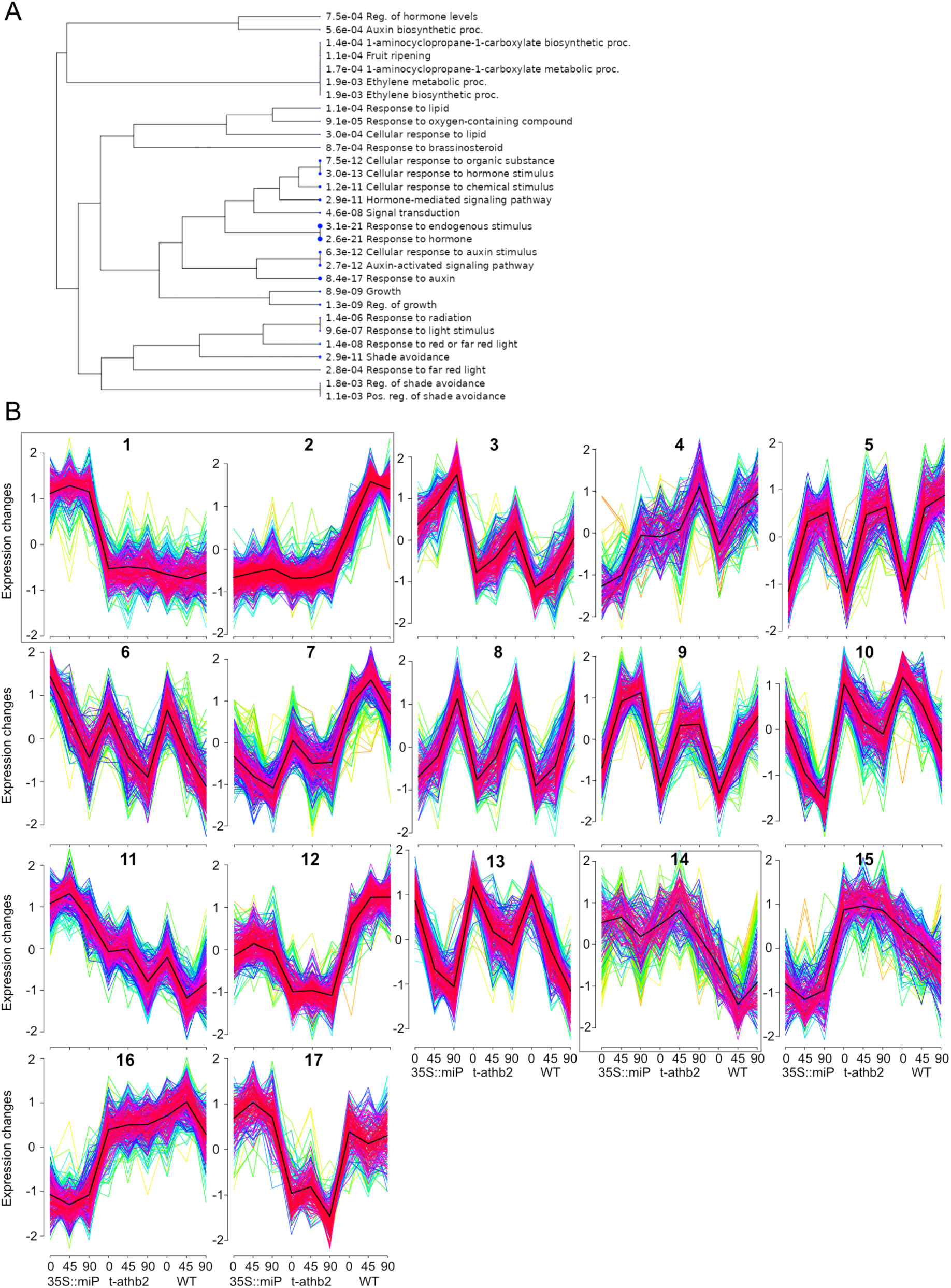
(A) GO enrichment analysis performed on the 95 DEGs whose expression is shown in Figure 3 yielded several significant GO terms (FDR < 0.05). These GO terms are related to shade avoidance, hormone/auxin responses, and growth. **(B)** The 17 clusters generated by MFuzz. The expression changes of the genes included in each cluster are on the y-axis. The three genotypes and shade conditions are on the x-axis.

**Figure S4.**
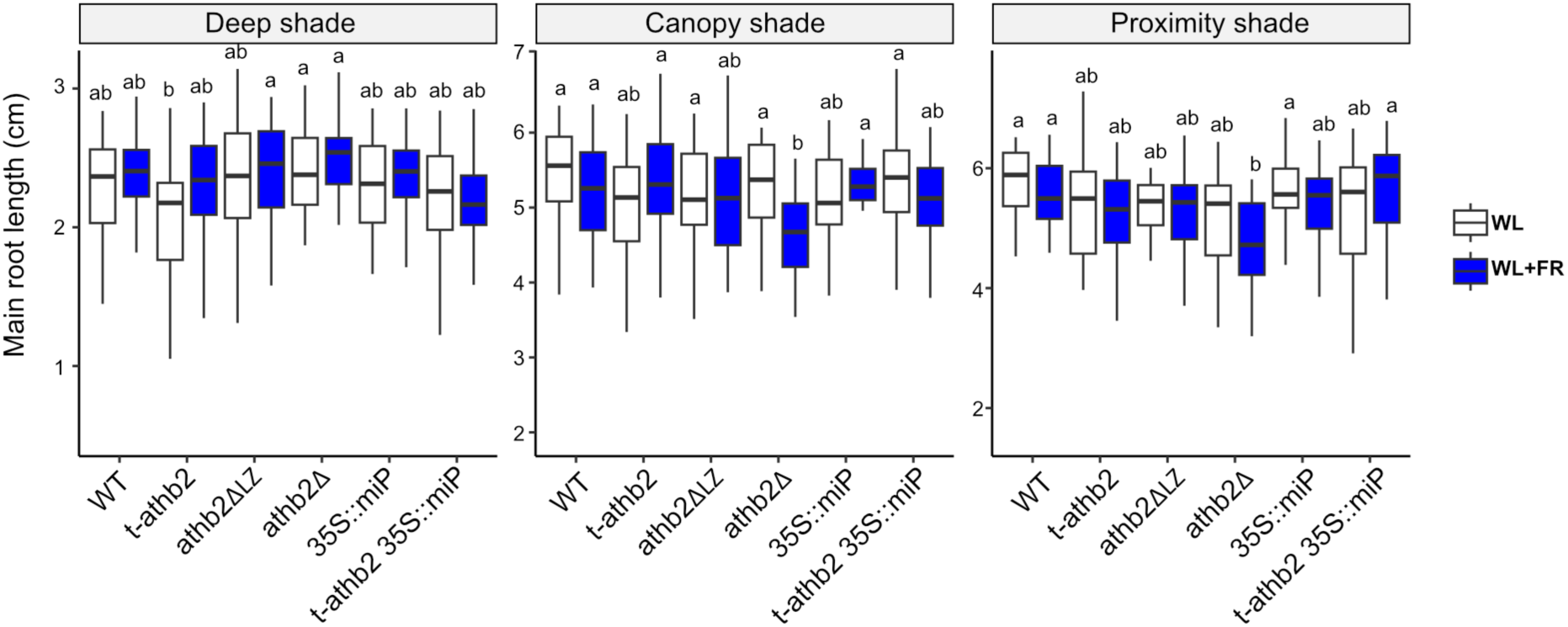
ATH2B and ATHB2miP slightly effect root elongation. Main root length of 11-day old plants grown in either WL or WL+FR in three different shade regimes: deep shade, canopy shade and proximity shade.

**Figure S5.**
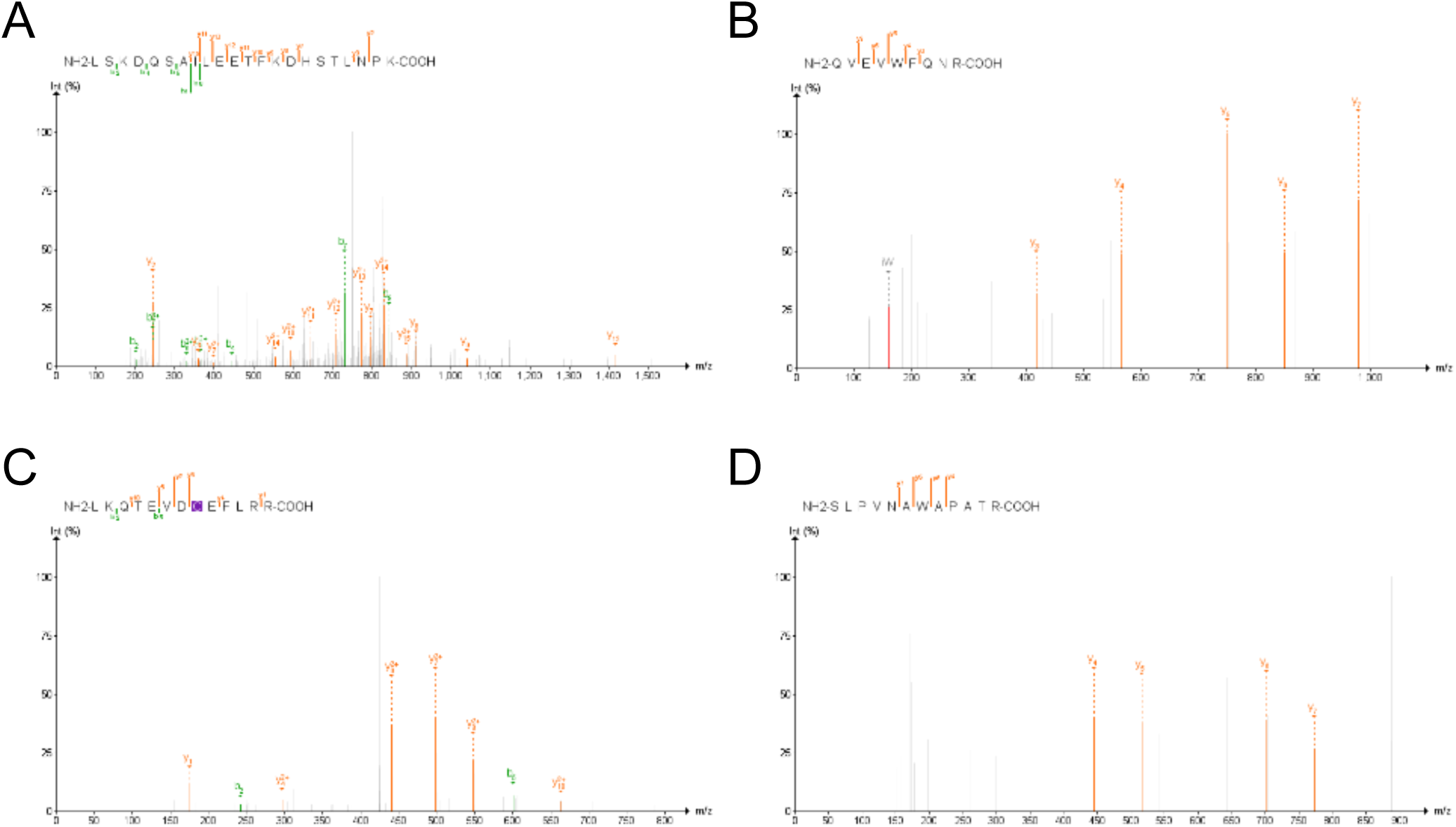
MS2 fragmentation spectra of ATHB2-derived peptides identified in complementation lines. (A-C) MS2 fragmentation spectra for peptides I, II, and III identified exclusively in ATHB2 WT complementation lines by immunoprecipitation mass spectrometry. (D) MS2 fragmentation spectrum for peptide IV identified exclusively in ATHB2 alt complementation lines. Peptide sequences are indicated above each panel.

## Notes

### Competing Interest Statement

The authors have declared no competing interest.

### Summary of Updates

The revised manuscript provides direct genetic and biochemical evidence for endogenous ATHB2miP function. A complementation experiment using a synonymous mutation at the ATHB2miP start codon demonstrates that loss of ATHB2miP production causes constitutively enhanced hypocotyl elongation, while protein analysis of the same lines confirms that this substitution alters ATHB2 proteoform composition. Interaction studies have been expanded with quantified FRET-FLIM data and EMSA experiments establishing the mechanistic basis for ATHB2miP activity. Multiple independent transgenic lines, corrected statistical analyses, and a full candidate dataset are also provided.

